# The Free Lunch is not over yet – Systematic Exploration of Numerical Thresholds in Phylogenetic Inference

**DOI:** 10.1101/2022.07.13.499893

**Authors:** Julia Haag, Lukas Hübner, Alexey M. Kozlov, Alexandros Stamatakis

## Abstract

Maximum Likelihood (ML) is a widely used phylogenetic inference model. ML implementations heavily rely on numerical optimization routines that use internal numerical thresholds to determine convergence. We systematically analyze the impact of these threshold settings on the log-likelihood and runtimes for ML tree inferences with RAxML-NG, IQ-TREE, and FastTree on empirical datasets. We provide empirical evidence that we can substantially accelerate tree inferences with RAxML-NG and IQ-TREE by changing the default values of two such numerical thresholds. At the same time, altering these settings does not significantly impact the quality of the inferred trees. We further show that increasing both thresholds accelerates the RAxML-NG bootstrap without influencing the resulting support values. For RAxML-NG, increasing the likelihood thresholds *ϵ*_LnL_ and *ϵ*_brlen_ to 10 and 10^3^ respectively results in an average tree inference speedup of 1.9 *±* 0.6 on *Data collection 1*, 1.8 *±* 1.1 on *Data collection 2*, and 1.9 *±* 0.8 on *Data collection 2* for the RAxML-NG bootstrap. Increasing the likelihood threshold *ϵ*_LnL_ to 10 in IQ-TREE results in an average tree inference speedup of 1.3 *±* 0.4 on *Data collection 1* and 1.3 *±* 0.9 on *Data collection 2*.

## Introduction

Phylogenetic trees have many important applications in biology and medicine, for example, in drug development (Gregoretti et al., 2004), forensics (Metzker et al., 2002), or the analysis of SARS-CoV-2 genomes (Morel et al., 2020). A widely used approach for reconstructing phylogenetic trees from a multiple sequence alignment (MSA) is the maximum likelihood (ML) method (Yang et al., 1995). Popular ML-based tools are RAxML-NG (Kozlov et al., 2019), IQ-TREE (Minh et al., 2020), and FastTree (Price et al., 2010). Finding the most likely tree is 𝒩𝒫-hard (Chor and Tuller, 2005) due to the super-exponential number of possible tree topologies. ML tree inference tools therefore implement tree search heuristics that attempt to iteratively optimize the log-likelihood (LnL score) by improving the tree topology, branch lengths, and substitution model parameters. These heuristics heavily rely on a plethora of numerical optimization routines (e.g., typically Brent’s method (Brent, 1971) and the Broyden-Fletcher-Goldfarb-Shanno (BFGS) method (Fletcher, 2000)) that use specific internal numerical convergence thresholds. To the best of our knowledge, the impact of these threshold settings on inference times and LnL scores has never been systematically assessed, while anecdotal observations do exist. For instance, when analyzing SARS-CoV-2 data, Morel et al. (2020) observed that one of these numerical thresholds, the minimum allowed branch length (*minBranchLen*), impacts the LnL scores of trees inferred with RAxML-NG and IQ-TREE. Here, we systematically investigate if we can reproduce this effect on other MSAs as well as for additional numerical thresholds. In addition to RAxML-NG and IQ-TREE, we also investigate the behavior of FastTree. We explore the influence of up to seven distinct numerical thresholds on LnL scores and runtimes for these three ML inference tools. For each tool, we analyze the influence of these settings on the standard tree inference procedure. During this standard tree inference procedure, the tools strive to optimize the tree topology, the branch lengths, and the substitution model parameters based on an initial starting tree topology. RAxML-NG and IQ-Tree also offer a tree evaluation procedure. During this tree evaluation procedure, the given (user-defined) tree topology remains fixed while only the branch lengths and substitution model parameters are being optimized. For RAxML-NG and IQ-Tree, we also analyze the influence of the numerical thresholds on this tree evaluation procedure.

A frequently used execution flavor in RAxML-NG is the --all mode. In addition to inferring 20 ML trees, RAxML-NG infers bootstrap replicate trees, and draws the bootstrap support values onto the best-scoring out of the 20 inferred ML trees. By default, RAxML-NG infers at most 1000 bootstrap replicates, but implements an early-stopping criterion that determines convergence based on the bootstopping criterion introduced by Pattengale et al. (2010). For RAxML-NG, we also analyze the influence of two likelihood epsilon thresholds on the results and the runtime of the bootstrapping procedure.

Our analyses comprise four main studies. The following paragraph summarizes each study and highlights the most important results.

### Study 1

In this first exploratory study, we analyzed the influence of up to 7 numerical thresholds on the LnL scores and runtimes of the RAxML-NG, IQ-TREE, and FastTree tree inference procedures. For RAxML-NG and IQ-TREE, we also analyzed the influence of the same thresholds when varied during the tree evaluation procedure. In this first study, we exclusively analyzed unpartitioned empirical DNA MSAs (*Data collection 1*). We observe a substantial runtime impact on tree inferences for two likelihood epsilons in RAxML-NG (*ϵ*_LnL_ and *ϵ*_brlen_). We further find that we can increase the settings of both thresholds without compromising the quality of the inferred trees, while obtaining a speedup of 1.9 *±* 0.6. We make a similar observation for one numerical threshold (*ϵ*_LnL_) in IQ-TREE that yields a speedup of 1.3 *±* 0.4. All other thresholds we analyzed for RAxML-NG and IQ-TREE, as well as all thresholds analyzed for FastTree do not substantially influence neither runtime nor LnL scores as long as these settings remain within a reasonable range. For all analyzed ML inference tools, their current default settings fall within this reasonable range. As expected, we observe that the runtime of the evaluation phase is small compared to the corresponding tree inference time. Despite the impact of some numerical thresholds on tree evaluation runtimes, we therefore recommend using a conservative numerical threshold setting for tree evaluation.

### Study 2

To verify the findings of *Study 1* for the likelihood epsilons *ϵ*_LnL_ and *ϵ*_brlen_ in RAxML-NG, and *ϵ*_LnL_ in IQ-TREE, we subsequently analyzed a more comprehensive as well as representative collection of empirical MSAs, including DNA, amino-acid (AA), and partitioned MSAs (*Data collection 2*). Our analyses on this more comprehensive data collection confirm our observations regarding tree inferences: the speedup for RAxML-NG is 1.8*±*1.1 and 1.3*±*0.9 for IQ-TREE. Analogous to our results on *Data collection 1*, we do not observe a significant impact on the quality of the inferred trees according to our evaluation metrics.

### Study 3

In our third study, based on the results of *Study 2*, we analyze the impact of the *ϵ*_LnL_ and *ϵ*_brlen_ thresholds on the RAxML-NG bootstrapping procedure. *Study 2* suggests that both thresholds can be increased for tree inferences without compromising the quality of the inferred trees, yet resulting in faster analyses. The hypothesis is that we can safely increase both thresholds to accelerate bootstrapping as well. We test this hypothesis using the MSAs of *Data collection 2*. Our analyses suggest that both likelihood epsilon settings can be increased without compromising the bootstrapping results and yield a speedup of 1.9 *±* 0.8 for *Data collection 2*.

### Study 4

In our final study, we conducted a more detailed analysis of the likelihood epsilons in RAxML-NG as it is being actively developed in our lab. Since RAxML-NG uses the same threshold *ϵ*_LnL_ for four distinct operations during its tree inference procedure, we separated this threshold into four distinct fine-grained likelihood epsilons. The goal was to assess if appropriate fine-grained threshold settings further improve runtimes. Our analyses suggest that separating the *ϵ*_LnL_ into four distinct thresholds does not further improve runtimes. We observe a similar behavior for all four thresholds, both in terms of tree inference quality and runtime. We hence conclude that such a fine-grained distinction of threshold settings is neither necessary nor beneficial. The remainder of this paper is organized as follows: In Methods, we outline the numerical thresholds we analyze and their usage in ML inference tools, our experimental setup, and the metrics we used to assess the influence of the numerical thresholds on tree inference quality, bootstrapping quality, and runtime. In Discussion we present our key findings and in Discussion we discuss the results of our analyses. To limit the extent of this paper, we only describe and discuss the results of *Study 2* and *Study 3* in greater detail. The results of *Study 1* and *Study 4* are available in the supplementary material. Finally, we conclude in Conclusion.

All MSAs we used for our analyses, as well as all results, are available for download at https://cme.h-its.org/exelixis/material/freeLunch_data.tar.gz. Our data generation scripts are available at https://github.com/tschuelia/ml-numerical-analysis.

## Methods

### Numerical Thresholds

Due to the extremely large tree space, an exhaustive search to identify the most likely tree is not feasible. ML-based tree inference tools therefore typically implement iterative tree improvement techniques, which they apply to an initial (starting) tree. Such an initial topology is obtained via heuristic tree inference methods (e.g., randomized stepwise addition order (Cavalli-Sforza and Edwards, 1967) or maximum parsimony (Farris, 1970; Fitch, 1971)). In our analyses, we focus on the three widely used ML inference tools RAxML-NG, IQ-TREE, and FastTree. Each tool iteratively optimizes the tree topology, the branch lengths, and the substitution model parameters starting from an initial tree. For example, RAxML-NG iteratively applies Subtree Pruning and Regrafting (SPR) moves followed by branch length and substitution model parameter optimizations. We provide a more detailed description of the tree search heuristics in Section 1 of the supplementary information. In our initial exploratory study *Study 1* we analyze the influence of the following seven numerical thresholds:

- Likelihood epsilon *ϵ*_LnL_: Threshold for LnL score improvement after one complete iteration (tree topology, branch lengths, and model parameters). The optimization only continues if the likelihood improvement is higher than this threshold.
- Branch length likelihood epsilon *ϵ*_brlen_: RAxML-NG specific threshold for LnL score improvement. This epsilon is used during a so-called fast branch length optimization to rapidly approximate the LnL score of potential SPR moves.
- Minimum branch length (*minBranchLen*): Lower limit for branch length values.
- Maximum branch length (*maxBranchLen*): Upper limit for branch length values.
- Model likelihood epsilon *ϵ*_model_: Threshold for substitution model parameter improvement. The substitution model parameters are only further optimized if the LnL score improvement exceeds this threshold.
- *num iters*: Threshold to control the maximum number of iterations during Newton-Raphson based branch length optimization in RAxML-NG.
- *bfgs factor* : This RAxML-NG specific threshold controls the convergence of the L-BFGS-B method used for optimizing substitution rates and stationary frequencies. The L-BFGS-B is a variant of the standard BFGS method, optimized for limited memory, and is extended to incorporate bound constraints in variables (Zhu et al., 1997).

For RAxML-NG we analyze the influence of all seven thresholds, for IQ-TREE we analyze the influence of *ϵ*_LnL_, *minBranchLen, maxBranchLen*, and *ϵ*_model_. For FastTree we analyze the influence of *ϵ*_LnL_ and *minBranchLen*. We provide a list of the threshold settings we analyze in Section 3 of the supplementary material. In all follow-up studies (*Studies 2–4*) we focus on the following thresholds: *ϵ*_LnL_ and *ϵ*_brlen_. In *Study 1* we find that decreasing the default settings does not substantially improve the LnL scores. To economize on computational resources and runtime, we thus only compare the current default setting to larger settings in *Studies 2–4*. For IQ-TREE, the current default setting for *ϵ*_LnL_ is 10^−3^. We analyze the potentially more liberal/superficial settings *{*10^−3^, 10^−2^, …, 10^3^*}*. For RAxML-NG the current default setting for both, *ϵ*_LnL_ and *ϵ*_brlen_, is 10^−1^. We analyze more superficial settings of *{*10^−1^, 1, …, 10^3^*}*

### Data Collections

In our exploratory *Study 1* we analyze 22 empirical unpartitioned DNA MSAs (*Data collection 1*). For all follow-up studies (*Studies 2–4*), we analyze a broader collection of 19 empirical MSAs, including AA and partitioned MSAs (*Data collection 2*). For one additional AA dataset with excessive memory and runtime requirements, we only compare the results of the default threshold settings to the suggested new default settings. We exclusively analyze empirical datasets, because it was shown that reconstructing the best tree is more difficult on empirical datasets than it is on simulated datasets (Huelsenbeck, 1995). Section 2 in the supplementary information provides a detailed overview of all MSAs we used for our analyses.

### Experimental Setup

In this section, we describe the experimental setup of our analyses. We separate this section into two paragraphs. In the first paragraph, we describe our experiments for analyzing the influence of the numerical thresholds on the tree inference and tree evaluation procedures (*Studies 1, 2, 4*). In the second paragraph, we describe our experiments for analyzing the influence of the likelihood epsilons on the bootstrapping procedure in RAxML-NG (*Study 3*). A more detailed description of our setup, as well as the software we use, is available in Section 1 of the supplementary information.

#### Tree Inference and Tree Evaluation

We analyze each threshold and each ML inference tool separately. For each threshold and for each possible threshold setting, we infer 50 trees using the standard/default tree inference mode of the respective tool. Subsequently, we re-evaluate each inferred tree using the tree evaluation mode. During the tree evaluation, we set the numerical thresholds to their corresponding default values. In the set comprising *all* inferred trees under *all* analyzed threshold settings, we determine the tree with the best LnL score (henceforth referred to as *best-known tree*) and compare it to all other trees using several distinct phylogenetic statistical significance tests. For reasons, we detail further below, we do not compare all trees at once, but always conduct a pairwise comparison of each tree with the best-known tree. We collect trees that pass *all* significance tests in a so-called plausible tree set (see Morel et al. (2020) for the introduction of the term). All trees in such a plausible tree set are not significantly worse than the best-known tree under all statistical significance tests.

To analyze the influence of the numerical thresholds on the RAxML-NG and IQ-Tree tree evaluation procedures, we use an analogous setup. Of the above described pipeline, we reuse the 50 trees inferred under the current default setting of the respective numerical threshold. For each possible threshold setting, we re-evaluate each of the 50 trees using the tree evaluation mode and the respective threshold setting. The subsequent plausible tree set analysis is analogous to the setup for the tree inference procedure described above.

#### Bootstrapping

In *Study 3* we exclusively analyze the influence of the likelihood epsilons on the RAxML-NG bootstrapping procedure. Since the bootstrapping procedure is computationally expensive, we refrain from testing all possible *ϵ*_LnL_ and *ϵ*_brlen_ settings in contrast to Studies 1, 2, and 4. Based on our findings in *Study 2* (see Discussion), we only compare the current default settings *ϵ*_LnL_ = 0.1 and *ϵ*_brlen_ = 0.1 to our suggested new settings *ϵ*_LnL_ = 10 and *ϵ*_brlen_ = 10^3^. To compare the bootstrap results under both settings, we first infer 20 ML trees using RAxML-NG’s standard tree inference procedure. Based on our findings of *Study 2* (see Discussion), we set the likelihood epsilons to the suggested new settings *ϵ*_LnL_ = 10 and *ϵ*_brlen_ = 10^3^ during this tree inference procedure. For both parameter configurations ((0.1, 0.1), (10, 10^3^)), we separately infer bootstrap replicates using RAxML-NG and map the bootstrap support values onto all 20 inferred ML trees.

#### Model of Evolution

For all experiments described above, we set the substitution model according to the following rules: For the unpartitioned DNA MSAs we use the general time reversible (GTR) model (Tavaré, 1986) of nucleotide substitution as it is the most flexible and general model of nucleotide substitution. To account for among site rate heterogeneity, we also use four discrete Γ rate categories. The AA equivalent of the GTR model is the GTR20 (or PROTGTR) model. However, this model for AA data is very parameter rich. In particular, on datasets with weak phylogenetic signal (see below) the corresponding parameter estimates might thus be unstable. Instead, we use the LG substitution model (Le and Gascuel, 2008) with four discrete Γ rate categories for unpartitioned AA MSAs. For partitioned MSAs, we use the partition file as provided alongside the MSA in the respective data source (see Supplementary material).

### Evaluation Metrics

#### Tree Inference and Tree Evaluation

In the following, we compare LnL scores in percent rather than via absolute LnL unit difference, since the datasets cover a broad range of absolute LnL values (LnL scores range between approximately −90 (D4) and −13 000 000 (D37)). Thus, as LnL scores are reported on a log scale, the observed effects are greater than the percentages might suggest. Therefore, we use two additional quality metrics: statistical significance tests and Robinson-Foulds distances (RF-Distances) (Robinson and Foulds, 1981) which we describe further below. For evaluating the runtimes of the tree inferences, we compare the runtime of each tree inference in relation to the average runtime under the respective default setting. We report all speedups as mean *±* standard deviation. Note that in our analyses we do not compare inferred trees, LnL scores, runtimes, or evaluation metrics across ML inference tools. All described analyses and evaluations metrics are applied separately and independently to each tool.

#### Significance Tests

In order to compare the trees inferred under different threshold settings, we use the statistical significance tests implemented in IQ-TREE. IQ-TREE implements the following significance tests: the Kishino-Hasegawa (KH) test (Kishino and Hasegawa, 1989) and the Shimodaira-Hasegawa (SH) test (Shimodaira and Hasegawa, 1999), both in their weighted and unweighted variants, the Approximately Unbiased (AU) test (Shimodaira, 2002), as well as the Expected Likelihood Weight (ELW) test (Strimmer and Rambaut, 2002). We use the default IQ-TREE settings for the number of resampling of estimated log-likelihoods (RELL) replicates (10 000) and the significance level (*α* = 0.05). We further denote a tree passing all statistical tests when compared to the best-known tree as being *plausible*. As described above, we collect all plausible trees per threshold setting in a plausible tree set. In subsequent analyses, we also use the number of plausible trees per setting, that is, the size of the respective plausible tree sets, as well as the number of unique plausible tree topologies (*N*_pl_) and their average pairwise RF-Distance (*RF*_pl_). Since the significance tests can be biased by the number of trees in the candidate set (Strimmer and Rambaut, 2002), we remove identical tree topologies from the set of inferred trees prior to applying the tests. Despite this tree set cleaning, we observed some unexpected behavior by the significance tests. First, the ELW test computes a c-ELW score (posterior weight) for each tree, sorts the trees according to this score and accepts trees as being not significantly different until the sum of c-ELW scores exceeds a predefined threshold. In our case, numerous trees in the inferred tree set have highly similar LnL scores despite their topologies being different. Yet, the c-ELW score for such trees is identical. Therefore, for trees that have a c-ELW that is close to exceeding the predefined significance threshold, only some trees with the exact same c-ELW score are accepted while the remaining ones are rejected. This leads to trees being rejected despite having identical LnL score as some accepted trees. Further, re-running the significance tests with the same trees but in a different order leads to a different subset of trees being accepted. Instead of re-estimating the substitution model parameters of each candidate tree, IQ-TREE uses a given best tree to optimize these parameters and uses them for all other trees. As stated above, numerous trees have identical LnL scores, and therefore choosing the best tree according to the LnL score is ambiguous. We observe that the results of the significance tests vary largely depending on what tree is passed as the best tree, despite identical LnL scores. We provide an example for both scenarios in the supplementary information. For the above reasons, instead of comparing all trees in the inferred tree set to each other at once, we only compare each inferred tree separately via all significance tests in a pairwise manner to the best-known tree. However, the c-ELW test is not intended for pairwise comparisons and only rejects one of the trees if the LnL scores deviate largely. Therefore, we also use the RF-Distance metric, which we describe in the following section.

#### RF-Distances

For the tree inference experiments, we fix the random seed to ensure that tree inferences always initiate their search on the same starting tree, despite using different numerical threshold settings. Therefore, we can directly compare tree topologies inferred under different numerical threshold settings that started on the same starting tree. We compare these trees in a pairwise manner via the relative RF-Distance. If the RF-Distance between two trees, for example, one inferred under *ϵ*_LnL_ = 10^−1^ and one inferred under *ϵ*_LnL_ = 10^3^ is 0.0, then the tree inference converged to the same topology despite the different *ϵ*_LnL_ setting. However, an RF-Distance *>* 0 does not necessarily indicate that the tree is worse. For example, the plausible tree set generally comprises multiple distinct tree topologies which are not distinguishable via statistical significance tests. Therefore, when using this metric, we further compare these RF-Distances to the average pairwise RF-Distance between all plausible trees inferred under the default numerical threshold setting per tool (*default plausible trees*). We further refer to this RF-Distance as *default RF-Distance*. This *default RF-Distance* provides a notion of how topologically scattered the plausible trees are under the default numerical threshold settings. The higher the *default RF-Distance* is, the more rugged the tree space will be. If the *default RF-Distance* is greater or equal to the RF-Distance between trees inferred under different numerical threshold settings, we assume that these differences are due to the ruggedness of the tree space rather than the trees being worse. Note that we do not compare branch lengths and substitution model parameters across settings, as they are not directly comparable for distinct tree topologies. For identical tree topologies, a subsequent tree evaluation after the tree inference is highly recommended and should be performed under conservative likelihood epsilon settings.

#### Bootstrapping

To determine the influence of the likelihood epsilon settings on the quality of the RAxML-NG bootstrapping procedure, we adopt an analogous quality assessment strategy as Stamatakis et al. (2008). As described in Section Experimental Setup, we infer 20 ML trees per MSA. For each of these 20 ML trees, we compare the bootstrap support values drawn onto those 20 trees based on bootstrap replicates inferred under the current default setting (*ϵ*_LnL_ = *ϵ*_brlen_ = 0.1) to the support values drawn onto the same 20 trees based on bootstrap replicates inferred under the suggested new setting *ϵ*_LnL_ = 10 and *ϵ*_brlen_ = 10^3^. Since we draw bootstrap support values on the same 20 ML trees, we can directly compare the values on a branch-by-branch basis using the Pearson correlation. This correlation only quantifies the relative similarity across bootstrap values. Hence, we also quantify the absolute difference between support values. To this end, we compute the pairwise absolute difference between support values under the old versus the suggested new setting across all branches of all 20 ML trees. Since RAxML-NG implements a bootstopping procedure (Pattengale et al., 2010), the number of bootstrap replicates may differ between likelihood epsilon configurations. We therefore also compare the number of computed replicates. We further summarize all bootstrap replicates per likelihood epsilon configuration in a consensus tree and compare the consensus trees between configurations using the RF-Distance. In the following analyses, we denote the consensus tree of all replicates inferred under *ϵ*_LnL_ = *ϵ*_brlen_ = 0.1 as *C*_default_ and the consensus for all replicates inferred under the suggested new settings (10, 10^3^) as *C*new.

#### Phylogenetic Signal

The properties of the MSA influence the phylogenetic inference (Stamatakis, 2011). The stronger the so-called *phylogenetic signal* in the data is, the easier the phylogenetic analysis will be. This phylogenetic signal provides a notion of how informative the data is about the underlying evolutionary process (Lemey et al., 2009). In our study, we use the sites-per-taxa ratio as a proxy for the phylogenetic signal. The sites-per-taxa ratio is computed by dividing the number of sites by the number of taxa in the MSA. In general, the higher the sites-per-taxa ratio of the MSA, the better the phylogenetic signal of the data will be. In the following analyses, we will refer to MSAs with a sites-per-taxa ratio ≥ 80 as *good phylogenetic signal*. We refer to MSAs with a lower sites-per-taxa ratio as MSA with an *intermediate* or *weak phylogenetic signal*.

## Discussion

In the following discussion, we focus on the analysis of the influence of *ϵ*_LnL_ and *ϵ*_brlen_ on the RAxML-NG and IQ-TREE tree inference (*Study 2*), as well as the influence of both thresholds on the RAxML-NG bootstrapping procedure (*Study 3*). All findings apply to *Data collection 2*. We discuss the less interesting results of *Study 1* and *Study 4* in detail in the supplementary material (Sections 4 and 6). The threshold with the highest impact on the runtimes of the RAxML-NG and IQ-Tree tree inference procedures is the likelihood epsilon *ϵ*_LnL_. We further observe a substantial impact of the branch length likelihood epsilon *ϵ*_brlen_ on the runtime of the RAxML-NG tree inference. Our analyses suggest that increasing these likelihood epsilon settings for RAxML-NG and IQ-TREE leads to equally good results, requiring less CPU time. The same observation holds true for the RAxML-NG bootstrapping procedure. All figures in the following section show the results summarized over all MSAs of *Data collection 2*. If not stated otherwise, we removed outliers using Tukey’s fences (Tukey, 1977) with *k* := 3 for all figures depicting a speedup for better visualization. For the sake of completeness, we provide comprehensive speedup figures including all outliers in Section 5 of the supplementary material. In all box plots, a dashed vertical line indicates the mean, and a solid vertical line the median value. In the following, we discuss our analysis results for *Study 2* and *Study 3* on *Data collection 2*. In the first paragraph, we focus on the influence of the likelihood epsilons on tree inferences with RAxML-NG and IQ-Tree (*Study 2*). In the second paragraph, we present our results for the RAxML-NG bootstrapping procedure (*Study 3*).

### Study 2: Tree Inference

#### RAxML-NG

With increasing *ϵ*_LnL_ threshold in RAxML-NG, we observe an expected decrease in LnL scores for higher settings. Especially for *ϵ*_LnL_ settings ≥ 10^2^ the LnL scores deteriorate noticeably (Figure 1a). This is reflected by the proportion of tree inferences yielding a tree that is included in the plausible tree set (henceforth called a plausible tree) as well. For the RAxML-NG default setting *ϵ*_LnL_ = 10^−1^ on average 85 % of tree inferences yield a plausible tree, for 10^3^ on average only 83 % yield a plausible tree. Averaged across all datasets, the *RF*_pl_ increases from 0.13 (10^−1^) to 0.16 (10^3^), and *N*_pl_ from 17.4 to 24.6. For all datasets (except D15) the RF-Distances between trees inferred under *ϵ*_LnL_ ≤ 10 compared to the default setting *ϵ*_LnL_ = 10^−1^ are smaller or equal to the *default RF-Distance*. However, for settings of 10^2^ and 10^3^ this is not the case. The average RF-Distances between trees inferred under these settings compared to the default setting are higher than the *default RF-Distance*. The topological differences among trees inferred under settings of 10^2^ and 10^3^ to trees inferred under the current default setting 10^−1^ can therefore not only be explained by the rugged tree space alone. This observation holds true even for datasets with a good phylogenetic signal. We conclude that for *ϵ*_LnL_ settings ≥ 10^2^ RAxML-NG infers worse trees than for settings below 10^2^. The runtimes of RAxML-NG tree inferences decrease with higher *ϵ*_LnL_ settings (Figure 1b). On average, tree inferences under *ϵ*_LnL_ = 10^3^ run approximately twice as fast as tree inferences under *ϵ*_LnL_ = 10^−1^.

**Fig. 1:**
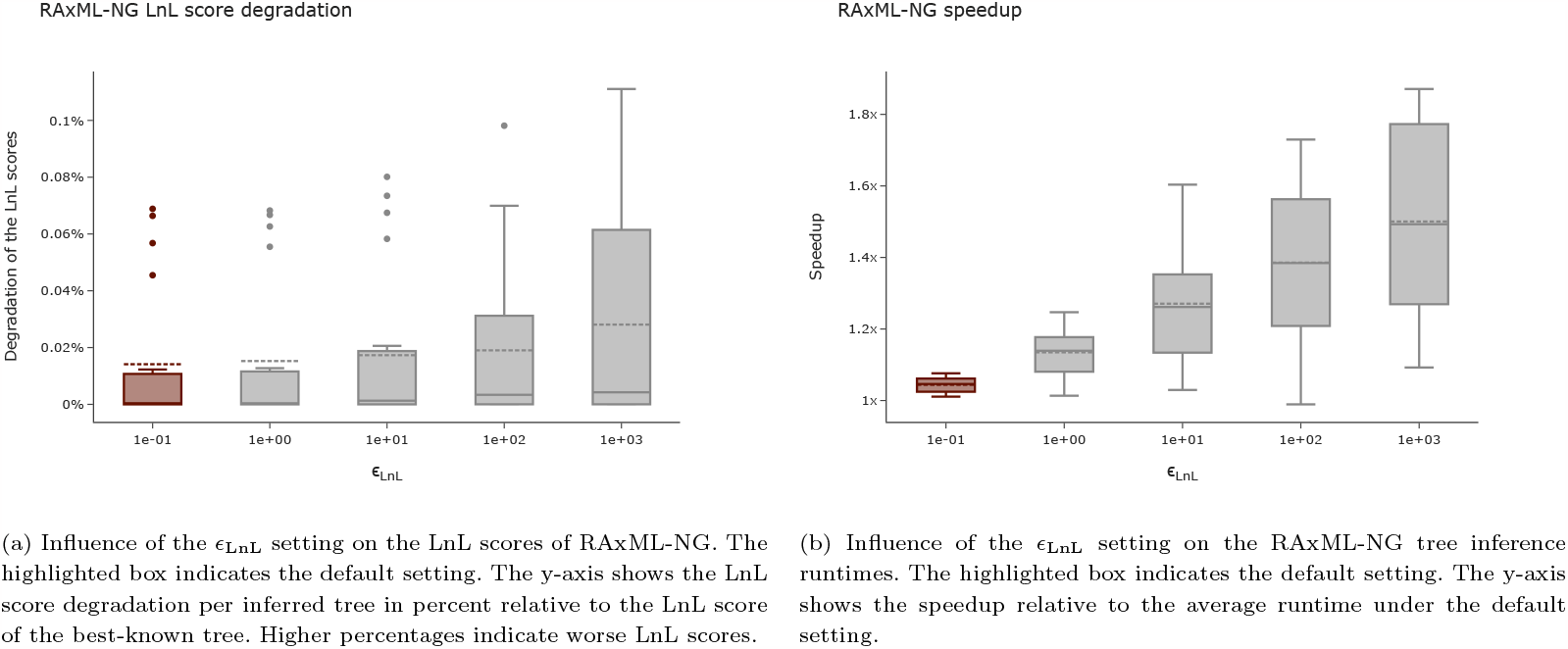
Influence of the *ϵ*_LnL_ setting on the LnL scores and runtime of RAxML-NG tree inferences.

Given these observations, we conclude that the *ϵ*_LnL_ setting can be increased to 10. The quality of the trees is not affected by this more superficial optimization, but the tree inferences run on average 1.4 *±* 0.6 times faster.

With RAxML-NG, we also analyze the influence of the *ϵ*_brlen_ threshold. Similar to the *ϵ*_LnL_ threshold, the runtimes for *ϵ*_brlen_ improve with increasing settings (Figure 2b). According to our analyses, the LnL score is unaffected by the *ϵ*_brlen_ setting (variations between settings ≤ 0.007 %; Figure 2a). Across all MSAs the number of tree inferences yielding a plausible tree is identical for all *ϵ*_brlen_ settings we analyze. The *RF*_pl_ increases only slightly from 0.13 (*ϵ*_brlen_ = 10^−1^) to 0.14 (*ϵ*_brlen_ = 10^−1^). In analogy, *N*_pl_ increases only slightly from 17.4 to 17.8 averaged over all datasets. For MSAs with a good phylogenetic signal, we observe that the *ϵ*_brlen_ setting does not affect the final tree topology: for all tested settings, the inferred tree topologies are identical (RF-Distance = 0.0). For all other MSAs, the average RF-Distance between trees inferred under different settings is below the *default RF-Distance*. We conclude that the *ϵ*_brlen_ threshold does not substantially influence the tree inference in RAxML-NG and the *ϵ*_brlen_ setting can be increased to 10^3^. In our analyses this observation holds true for all analyzed MSAs independently of the magnitude of the LnL scores. RAxML-NG uses the *ϵ*_brlen_ to optimize the three branch lengths that are adjacent to the node at which a subtree is regrafted via an SPR move. We suspect that since all branch lengths are optimized at a later step during the tree inference, conducting a thorough optimization of these three branch lengths does not substantially improve the LnL score and can thus be terminated early.

**Fig. 2:**
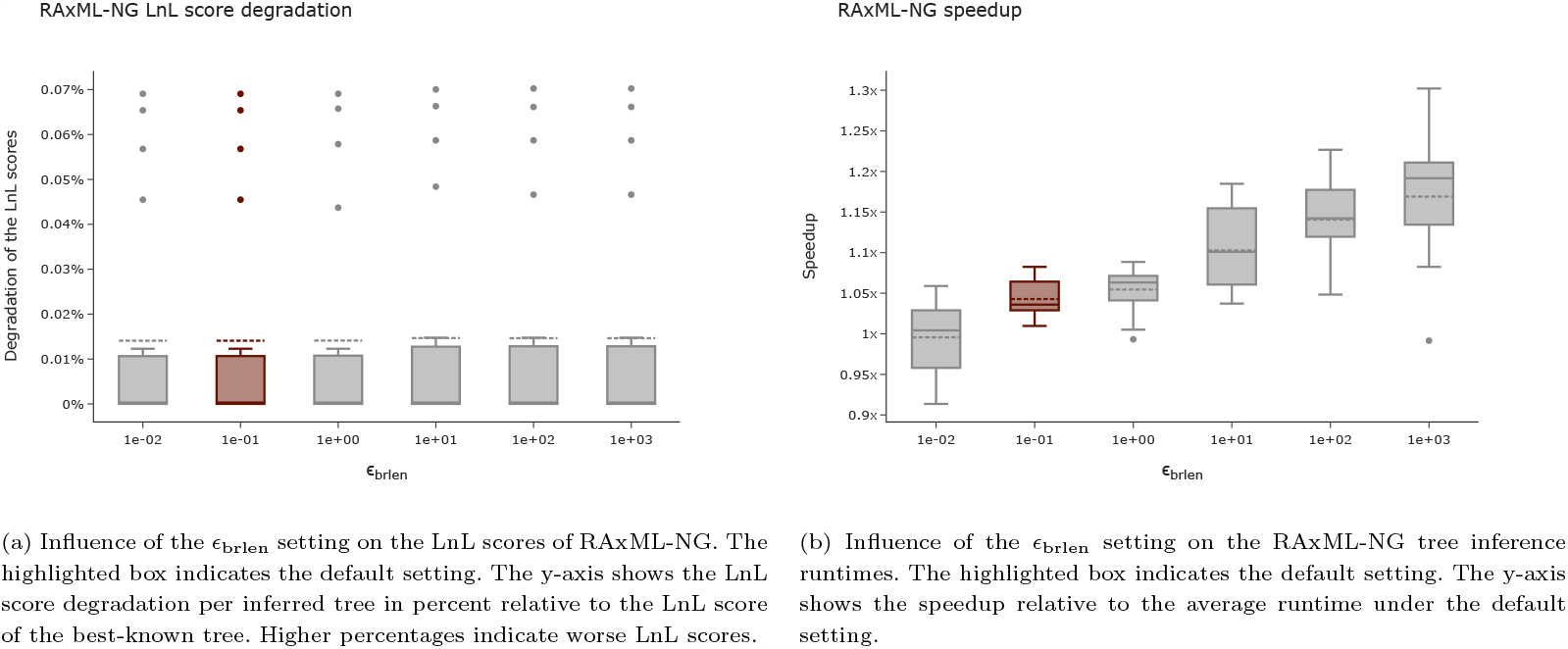
Influence of the *ϵ*_brlen_ setting on the LnL scores and runtime of RAxML-NG tree inferences.

Since we suggest changing two likelihood epsilons in RAxML-NG, we further analyze the influence of simultaneously changing both settings on the quality and the runtimes of tree inferences. To limit the computational effort, we only compare the default combination (*ϵ*_LnL_, *ϵ*_brlen_) = (10^−1^, 10^−1^) with the suggested new combination (*ϵ*_LnL_, *ϵ*_brlen_) = (10, 10^3^). As expected, the LnL scores are worse under the new setting compared to the old setting (Figure 3a), but the tree inferences are faster (Figure 3b). Averaged over all MSAs, the LnL scores between the current default and the suggested new combination vary by less than 0.004 %. The percentage of tree inferences yielding a plausible tree is identical under both setting combinations (87 %). We observe only a minor increase of *RF*_pl_ from 0.12 to 0.14 and *N*_pl_ from 17.2 to 21.4. For all MSAs the RF-Distances between trees inferred under the current default combination versus the new combination are smaller or equal to the *default RF-Distance*. We conclude that increasing both threshold settings does not substantially decrease the LnL scores of the inferred trees and does therefore not affect the quality of the inferred trees. With the MSAs of *Dataset collection 1* we observe a speedup of 1.9 *±* 0.6, on *Data collection 2* we observe a speedup of 1.8 *±* 1.1.

**Fig. 3:**
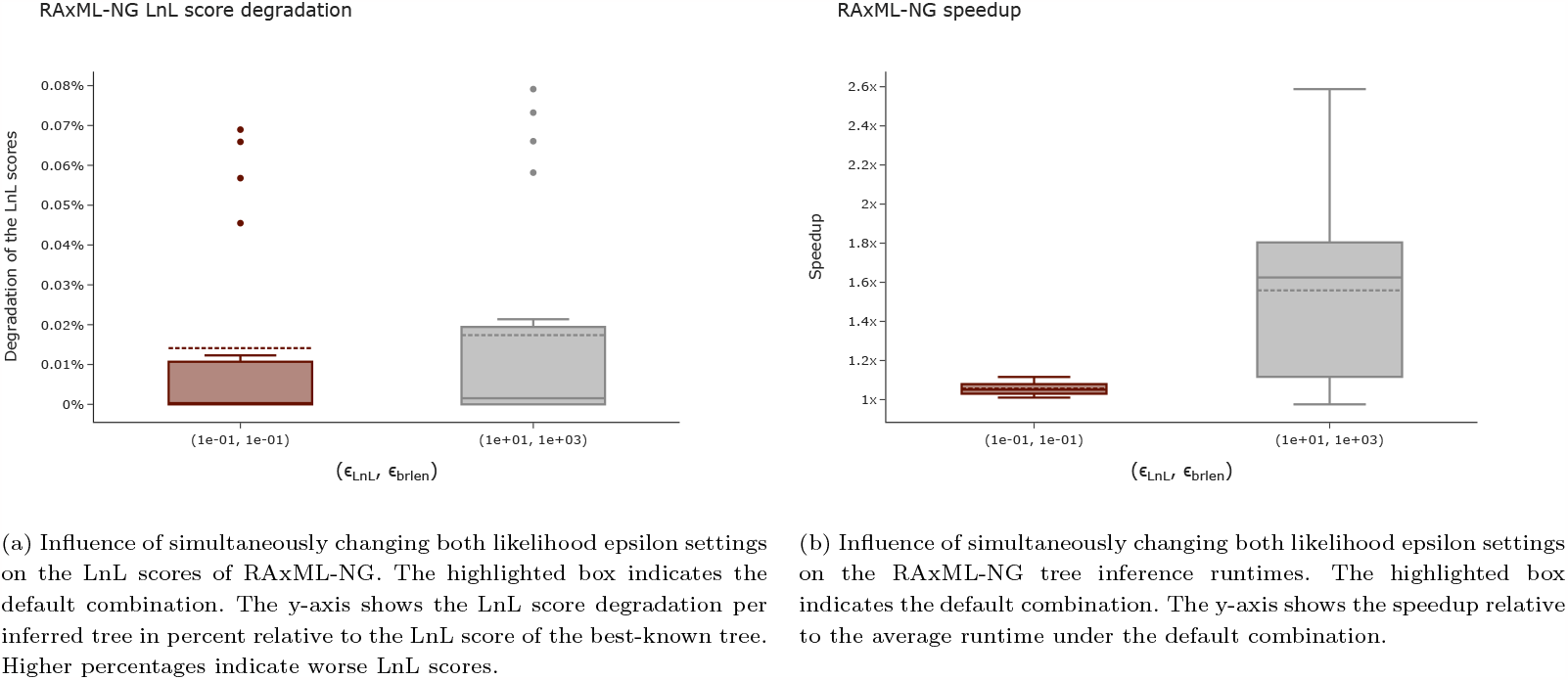
Influence of simultaneously changing both likelihood epsilon settings on the LnL scores and runtime of the RAxML-NG tree inference.

#### IQ-TREE

Analogous to RAxML-NG, the runtimes of tree inferences improve with higher *ϵ*_LnL_ settings for IQ-Tree. Tree searches under the default setting of *ϵ*_LnL_ = 10^−3^ run on average approximately twice as long as tree searches with *ϵ*_LnL_ = 10^3^ (Figure 4b). However, IQ-TREE appears to be more sensitive to the *ϵ*_LnL_ setting than RAxML-NG in terms of LnL scores. Under higher *ϵ*_LnL_ settings, the LnL score degradation is an order of magnitude worse than for RAxML-NG (on average ≤ 0.2 % for IQ-TREE vs. ≤ 0.03 % for RAxML-NG; Figure 4a). For *ϵ*_LnL_ values ≤ 10 the LnL scores are on average approximately equal. Also, based on the plausible tree set size under various settings, we observe that IQ-TREE is more sensitive to the *ϵ*_LnL_ setting. We observe that for *ϵ*_LnL_ = 10^3^ averaged over all MSAs, noticeably fewer tree inferences yield a plausible tree than for any other setting (58 % vs. 76 % for *ϵ*_LnL_ = 10^−3^). This effect is less pronounced for MSAs with good phylogenetic signal. For MSAs with a sites-per-taxa ratio ≥ 80 we observe that the *ϵ*_LnL_ setting does not affect the final tree topology: under all tested settings the inferred tree topologies are identical (RF-Distance = 0.0). For MSAs with a worse phylogenetic signal, the RF-Distance between trees inferred under the default setting 10^−3^ and settings of 10^2^ and 10^3^ exceed the average RF-Distance in the plausible tree set. We conclude that for MSAs with an intermediate or weak phylogenetic signal, the trees inferred under *ϵ*_LnL_ settings ≥ 10^2^ are worse than under lower settings. According to our evaluation metrics across all analyzed MSAs the *ϵ*_LnL_ setting can be set to 10 without compromising the quality of the inferred trees. In our analyses, this results in an average speedup of 1.3 *±* 0.9.

**Fig. 4:**
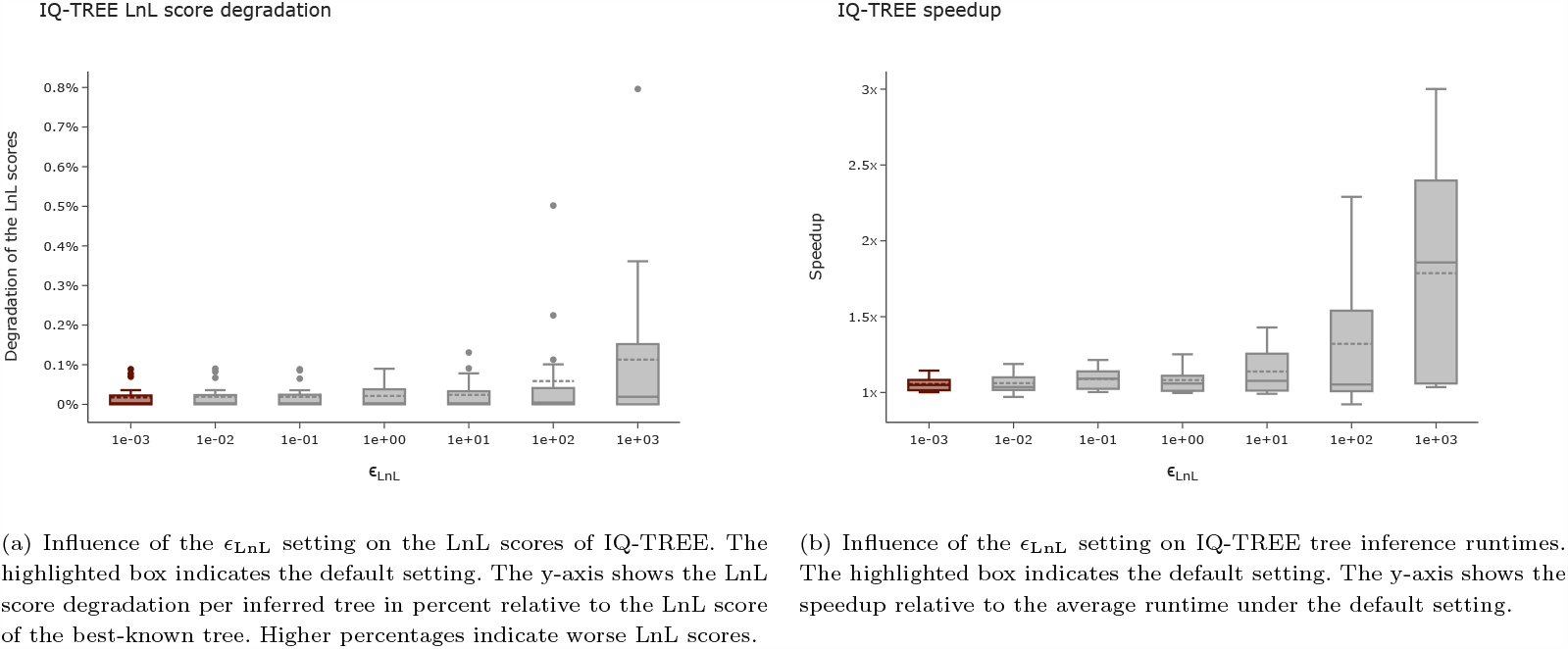
Influence of the *ϵ*_LnL_ setting on the LnL scores and runtimes of IQ-TREE tree inferences.

As mentioned before, we observe a higher sensitivity to the *ϵ*_LnL_ setting in IQ-TREE than in RAxML-NG. We suspect that this is caused by the random Nearest Neighbor Interchange (NNI) topology perturbation moves in IQ-TREE’s search algorithm. IQ-TREE implements these random NNI moves to escape local NNI maxima (see the supplementary information for a more detailed description of the IQ-TREE inference heuristic). To explore this hypothesis, we modify IQ-TREE and disable this randomness in the search algorithm. As a consequence, IQ-TREE then only optimizes the tree topology using standard NNI moves. We refer to the standard IQ-TREE as *random IQ-TREE* and to the IQ-TREE algorithm without random NNI moves as *de-randomized IQ-TREE*. We re-analyze four MSAs using the de-randomized IQ-TREE version. Without the random NNI moves, the IQ-TREE search heuristic can explore the tree space less, thus, we expect the LnL scores for de-randomized IQ-TREE to be worse than for random IQ-TREE, which we indeed observe in our analyses. To compare the influence of the *ϵ*_LnL_ threshold, we again compute the proportion of tree inferences yielding a plausible tree. We observed that when using de-randomized IQ-TREE, noticeably more tree inferences yield a plausible tree under *ϵ*_LnL_ ≥ 10^2^ than when using the random IQ-TREE variant. We conclude that large *ϵ*_LnL_ settings (≥ 10^2^) distort the random NNI moves in IQ-TREE, causing a premature termination of the tree inference. This also explains the vast runtime improvement under these settings.

### Study 3: RAxML-NG Bootstrapping

Note that in the following discussion, we will refrain from reporting average correlation coefficients, as the most accurate method of averaging correlations is disputed (Corey et al., 1998). Especially, the widely used method of applying the Fisher z-transformation (Fisher, 1992) prior to averaging is not directly applicable to our analyses results, as the inverse hyperbolic tangent function is only defined for values *<* 1.0. We do, however, frequently observe Pearson correlation coefficients of *exactly* 1.0.

We observe Pearson correlation coefficients *>* 0.99 for all 20 ML trees when comparing the support values under the current default setting *ϵ*_LnL_ = *ϵ*_brlen_ := 0.1 to the suggested new setting *ϵ*_LnL_ := 10 and *ϵ*_brlen_ := 10^3^ (Figure 5a). All p-values are ≤ 10^−35^. We observe the lowest correlation coefficient on D354 (0.992). This is, however, not surprising, as this dataset contains highly similar ITS sequences, and is known to be difficulty to analyze (Grimm et al., 2006). RAxML-NG reports support values in percent on a scale of 0% to 100%. Averaged over all datasets, the absolute pairwise difference in support values per tree topology is 0.5 percentage points. The highest observed difference is 3.23 percentage points for a phylogeny inferred on D25. On D25, the average support value over all 20 ML trees is 55.2%. This suggests that the bootstrap replicates inferred under both settings do not substantially influence the interpretation of the resulting support values drawn on the ML trees. As mentioned above, due to the implemented bootstopping procedure, the number of bootstrap replicates may differ depending on the likelihood epsilon setting. Except for two datasets, the computed number of bootstrap replicates is identical. For D354, we observe a slower convergence with the suggested new settings: RAxML-NG infers 700 replicates under (0.1, 0.1) and 800 under (10, 10^3^). In contrast, for D140, we observe a faster convergence with 400 inferred replicates under (0.1, 0.1) versus 350 replicates under (10, 10^3^). To further analyze this observation for D140 and D354, we reran the analyses using distinct random seeds. For D354, changing the random starting seed to 42 and 100 in distinct analyses, showed a similar trend: for seed = 42 the more conservative setting (0.1, 0.1) converges after 850 bootstrap replicates while it does not converge under (10, 10^3^) and infers the maximum number of 1000 replicates. With seed = 100 we observe 700 replicates under (0.1, 0.1) versus 900 replicates under (10, 10^3^). Yet, despite the increased number of inferred replicates, the increased likelihood epsilon settings result in a speedup *>* 1 compared to (0.1, 0.1). For D140, changing the random starting seed from 0 to 42 resulted in identical number of replicates under both likelihood epsilon configurations (384 replicates).

**Fig. 5:**
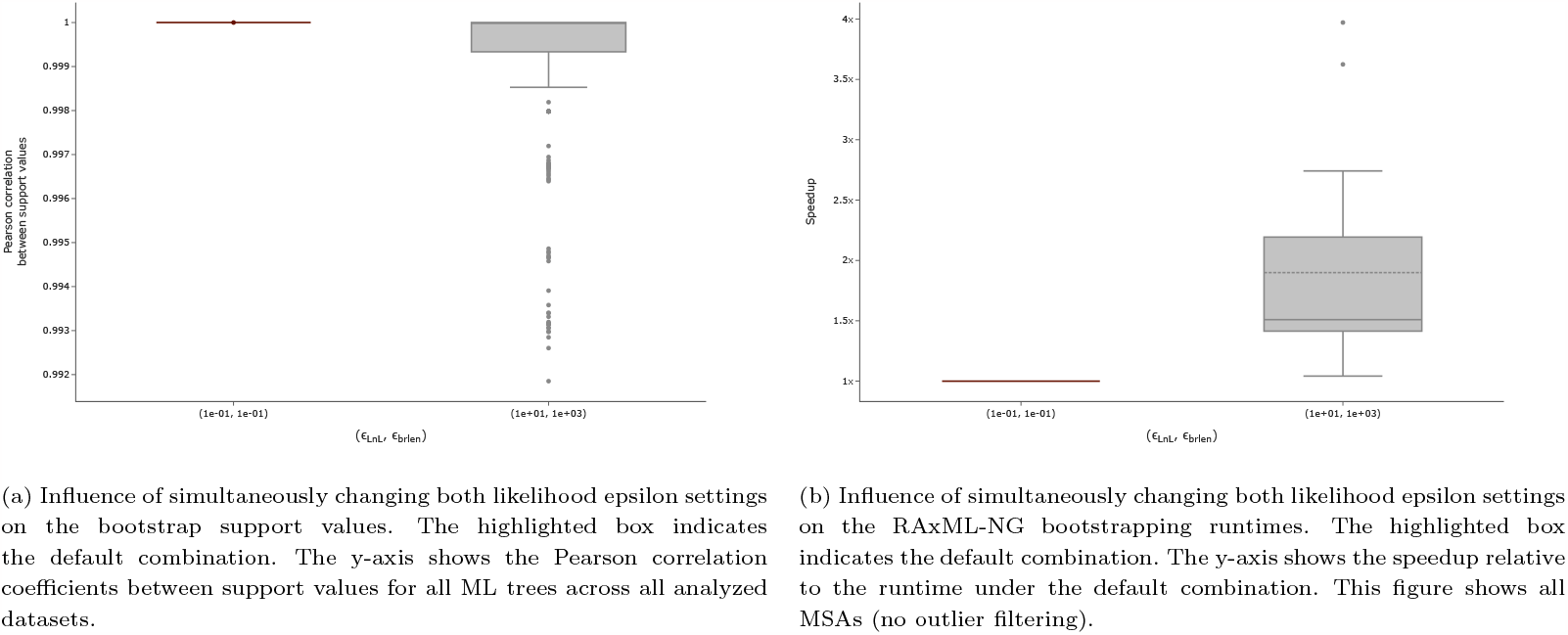
Influence of simultaneously changing both likelihood epsilon settings on the bootstrap support values and runtime of the RAxML-NG bootstrap.

Averaged over all MSAs, the relative RF-Distance between the respective bootstrap consensus trees *C*_default_ and *C*_new_ is 0.03. For 11 out of 20 MSAs, the consensus trees *C*_default_ and *C*_new_ are topologically identical (RF-Distance = 0.0). We observe the highest topological difference again for D354 (RF-Distance = 0.14). We conclude that the bootstrapping procedure is not affected by the increased likelihood epsilon values, and we suggest changing the respective default settings in RAxML-NG. Implementing the suggested changes results in a speedup of 1.9 *±* 0.8 on *Data collection 2* (Figure 5b).

## Conclusion

Increasing the RAxML-NG settings for the likelihood epsilons *ϵ*_LnL_ and *ϵ*_brlen_ to 10 and 10^3^ respectively does not significantly influence the quality of the inferred trees according to statistical significance tests. By changing both settings, we observe a speedup of 1.9 *±* 0.6 on *Data collection 1* and 1.8 *±* 1.1 on *Data collection 2*. With IQ-TREE, increasing the *ϵ*_LnL_ to 10 has no significant impact on the LnL scores, and we observe a speedup of 1.3 *±* 0.4 on *Data collection 1* and 1.3 *±* 0.9 on *Data collection 2*. Our observations are independent of the magnitude of the LnL scores of the analyzed MSAs. For MSAs with a good phylogenetic signal, the inferred tree topologies under the current default settings and the suggested new settings are identical for both, RAxML-NG and IQ-TREE (RF-Distance = 0.0). For MSAs with an intermediate or weak phylogenetic signal, the topological differences between threshold settings can be explained by the rugged tree space, and the RF-Distances between inferred trees under different settings are less than or equal to the *default RF-Distance*. It is important to note that the final tree evaluation after tree inference should not be omitted and performed under conservative likelihood epsilon settings, for example the default settings in RAxML-NG and IQ-TREE.

We further suggest increasing the *ϵ*_LnL_ and *ϵ*_brlen_ to 10 and 10^3^ respectively during the RAxML-NG bootstrapping procedure as well. According to our analyses, these changes do not affect the quality of the bootstrapping results while decreasing the runtime of the RAxML-NG bootstrap. We observe a speedup of 1.9 *±* 0.8 on *Data Collection 2*.

Based on our results, the default values for *ϵ*_LnL_ and *ϵ*_brlen_ were increased in the production level release of RAxML-NG Version 1.2.0 to 10 and 10^3^ respectively. RAxML-NG Version 1.2.0 further performs the suggested tree evaluation step with conservative likelihood epsilon settings (i.e. 0.1) after each tree inference (see https://github.com/amkozlov/raxml-ng/releases/tag/1.2.0).

## Funding

This work was financially supported by the Klaus Tschira Foundation, by a grant from the Ministry of Science, Research and the Arts of Baden-Württemberg (Az: 33-7533.-9-10/20/2) to Peter Sanders and Alexandros Stamatakis, and by the European Union (EU) under Grant Agreement No 101087081 (Comp-Biodiv-GR).

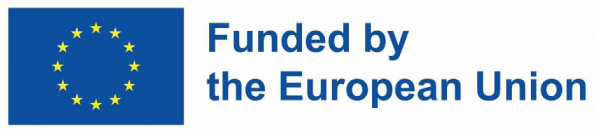

