## Supplementary material for "The Free Lunch is not over yet – Systematic Exploration of Numerical Thresholds in Phylogenetic Inference"

June 15, 2023

#### 1 Experimental Setup

In this section, we describe our experimental setup. First, we explain the tree inference heuristics of RAxML-NG, IQ-TREE, and FastTree in more detail. Afterwards, we provide the command lines we use for our analyses, as well as the respective software versions of the ML inference tools. Finally, we describe our pipeline setup and the hardware we perform our analyses on.

##### 1.1 ML Inference Tools

**RAxML-NG** RAxML-NG [10] was introduced in 2019 as a successor of the widely used ML inference tool RAxML [20]. RAxML-NG optimizes the initial starting tree using a greedy hill-climbing algorithm. Here, greedy means that only steps improving the LnL score are being accepted. The hill-climbing procedure comprises multiple rounds of optimizing the substitution model parameters, the branch lengths, and the tree topology. The branch lengths and substitution model parameters are optimized using the Newton-Raphson, L-BFGS-B [4], and Brent [1] methods. RAxML-NG allows the user to constrain the branch length values via the *minBranchLen* and *maxBranchLen* thresholds. The user can also define the convergence limit of the L-BGFS-B method via the threshold *bfgs_factor*.

RAxML-NG implements Subtree Pruning and Regrafting (SPR) moves as topology optimization strategy. One iteration of pruning and regrafting all possible subtrees in the current tree is called an *SPR round*. Instead of performing all possible reattachments, RAxML-NG only reattaches a pruned subtree into all neighboring branches up to a maximum distance of nodes away from the pruning position, the *SPR radius*. RAxML-NG automatically determines this SPR radius by performing SPR rounds and increasing the SPR radius until the LnL score does not further improve (*autodetect* SPR rounds). On the initial starting tree topology, RAxML-NG first performs so-called *fast* SPR rounds. During these fast SPR rounds, the branch lengths around subtree regraft nodes are not optimized prior to scoring the tree. During the subsequent *slow* SPR rounds, the lengths of the three branches adjacent to the regrafting node are optimized prior to computing the LnL score. After each fast or slow SPR round, RAxML-NG optimizes all branch lengths of the 20 most promising tree topologies found during this SPR round. Once the LnL score does not improve more than by $\epsilon_{\text{LnL}}$, RAxML-NG terminates the SPR rounds. Before exiting, RAxML-NG optimizes the substitution model parameters until the LnL score converges, that is, if the LnL score does not improve by at least $\epsilon_{\text{model}}$ between model parameter optimization iterations.

When optimizing all branch lengths, RAxML-NG repeatedly iterates over all branches in the tree to improve their values with respect to the LnL score. The user can limit the maximum number of passes over the entire tree via the parameter *num_iters*. Throughout the tree inference process, RAxML-NG uses likelihood epsilons to detect LnL score convergence during these numerical optimization procedures. After each iteration, RAxML-NG checks if the likelihood improved by at least a certain threshold. Standard RAxML-NG uses two different likelihood epsilons: $\epsilon_{\text{LnL}}$ and $\epsilon_{\text{brlen}}$. RAxML-NG uses the $\epsilon_{\text{LnL}}$ threshold during the autodetect, fast, and slow SPR rounds, as well as when it optimizes all branch lengths in the tree. For our more thorough likelihood epsilon study *Study 4*, we modified RAxML-NG such that it uses a dedicated threshold for each of these four steps. The $\epsilon_{\text{brlen}}$ threshold is used during the slow SPR rounds, when RAxML-NG optimizes the three branch lengths adjacent to the pruning node. In Study 4, we distinguish between the following likelihood epsilons:

- $\epsilon_{\text{auto}}$: RAxML-NG uses this threshold during the autodetect SPR rounds. The SPR radius is increased and RAxML-NG continues running autodetect SPR rounds, if the LnL improved by at least $\epsilon_{\text{auto}}$.
- $\epsilon_{\text{fast}}$: RAxML-NG uses this threshold during the fast SPR rounds. RAxML-NG continues performing fast SPR rounds, as long as the LnL score improves by at least $\epsilon_{\text{fast}}$.
- $\epsilon_{\text{slow}}$: RAxML-NG uses this threshold during the slow SPR rounds. RAxML-NG continues performing slow SPR rounds, as long as the LnL score improves by at least $\epsilon_{\text{slow}}$.
- $\epsilon_{\text{brlen\_full}}$ RAxML-NG uses this threshold when it optimizes all branch lengths.
- $\epsilon_{\text{brlen}}$ RAxML-NG uses this threshold during the slow SPR moves when it optimizes the branch lengths of the three adjacent branches to decide when to stop iterating over these three branches for adjusting their values.

The standard RAxML-NG implementation summarizes the thresholds $\epsilon_{\text{auto}}$, $\epsilon_{\text{fast}}$, $\epsilon_{\text{slow}}$, and $\epsilon_{\text{brlen\_full}}$ as the single threshold $\epsilon_{\text{LnL}}$. Since the threshold $\epsilon_{\text{brlen}}$ is only used at one step in the tree inference process, this threshold remains unchanged.

**IQ-TREE** IQ-TREE [13] is a widely used software package for phylogenetics that was first released in 2014. Similar to RAxML-NG, IQ-TREE in principle also implements a greedy hill-climbing ML tree inference. IQ-TREE's topology optimization strategy consists in repeated NNI moves. Since NNI moves explore the tree space less exhaustively than SPR moves [12], IQ-TREE also implements random moves to conduct backward steps with respect to the LnL score. This allows IQ-TREE to explore the tree space beyond the NNI neighborhood and navigate out of local maxima. These random NNI moves are applied to a locally optimal tree $T_{\text{best}}$ with LnL score $\text{LnL}_{\text{best}}$. A random NNI move perturbs the tree by applying a randomly selected NNI move. After this random NNI move, IQ-TREE performs repeated standard NNI moves until the tree is NNI optimal, meaning that any additional NNI move will decreases the LnL score again. If this new tree $T^*$ has a higher LnL score than $\text{LnL}_{\text{best}}$, IQ-TREE replaces $T_{\text{best}}$ with $T^*$ and repeats the search cycle. If the LnL score of $T^*$ is lower than $\text{LnL}_{\text{best}}$, IQ-TREE discards $T^*$ and repeats the search cycle using the current $T_{\text{best}}$. Since the random NNI move is selected at random, the repeated cycle will, with high probability, select a different random NNI move. During the repeated standard NNI moves, IQ-TREE additionally optimizes the branch lengths until the LnL score increases by less than $\epsilon_{\text{LnL}}$. This entire cycle of random NNI moves and standard NNI moves is repeated until IQ-TREE does not find a better tree than $T_{\text{best}}$ during 100 cycles. Before exiting, IQ-TREE implements a final optimization of substitution model parameters and checks for convergence of this last optimization step using the $\epsilon_{\text{LnL}}$ threshold. Similar to RAxML-NG, IQ-TREE also optimizes branch lengths and model parameters using the Newton-Raphson, L-BFGS-B, and Brent methods. One can constrain the branch length value range via the *minBranchLen* and *maxBranchLen* thresholds.

**FastTree** FastTree [14] aims to reduce the time and space complexity of ML phylogenetic inference. The authors achieve this through various heuristics and shortcuts compared to other, more standard, ML inference tools. In contrast to RAxML-NG and IQ-TREE, FastTree initially conducts a minimum evolution criterion [28] based inference step before maximizing the tree's likelihood. The minimum evolution approach attempts to obtain a tree that explains the data with as few mutations as possible, therefore minimizing the branch lengths, with the branch lengths being at least *minBranchLen*. The minimum evolution criterion search steps include NNI and SPR moves. Instead of iterating until convergence, FastTree runs a predefined number of minimum evolution search rounds. For the subsequent ML optimization, FastTree uses NNI steps to improve the tree topology with respect to its LnL score. FastTree executes at most $2 \cdot N$ NNI rounds, where $N$ is the number of taxa in the MSA. During these NNI rounds, FastTree compares the current LnL score of the tree to the previous LnL score and terminates the NNI rounds if it increases by less than $\epsilon_{\text{LnL}}$.

| Tool | Software Version |
| --- | --- |
| RAxML-NG Study1 | Adapted version that allows setting of additional numerical thresholds via the command line. Based on version 1.0.1. Available at `https://github.com/lukashuebner/param-raxml-ng` |
| RAxML-NG *Study 2* | Adapted version that separates the $\epsilon_{\text{LnL}}$ into four distinct thresholds, where each can be set separately via the command line. Based on version 1.1.0. Available at `https://github.com/tschuelia/raxml-ng`. |
| IQ-TREE | Version 2.1.3, available at `https://github.com/iqtree/iqtree2/releases/tag/v2.1.3` |
| de-randomzied IQ-Tree | Adapted IQ-TREE version that does not perform random NNI moves. Based on version 2.1.3. Available at `https://github.com/tschuelia/derandomized-iqtree` |
| FastTree | Adapted version that allows to set *minBranchLen* and $\epsilon_{\text{LnL}}$ via the command line. Based on version 2.1. Available at `https://github.com/tschuelia/param-fasttree` |

Table S1: The software versions of RAxML-NG, IQ-TREE, FastTree, and RAxML.

#### 1.2 Software and Command Lines

In our analyses we use RAxML-NG, IQ-TREE, and FastTree to infer phylogenetic trees. We analyze the influence of the numerical thresholds for each tool separately and do not compare results across different ML inference tools. With RAxML-NG, we re-evaluate the inferred trees using its own tree evaluation mode. Analogously, we re-evaluate trees inferred with IQ-TREE using the IQ-TREE tree evaluation mode. Since FastTree does not provide a tree evaluation mode, we do not re-evaluate trees inferred using FastTree. For all three ML inference tools, we use the significance tests implemented in IQ-TREE to determine the plausible tree sets. We infer bootstrap replicates and draw support values using RAxML-NG. We perform the bipartition frequency correlation analyses using RAxML. Table S1 states the software versions we use, and Table S2 shows the command lines used for the respective task and tree inference tool.

| Mode | Tool | Command Line |
|---|---|---|
| Search | RAxML-NG *Study 1* | `raxml-ng --msa` $\langle$MSA$\rangle$ `--model` $\langle$model/partition$\rangle$ `--tree` $\langle$pars{1}/rand{1}$\rangle$ `--blmin` $\langle$*minBranchLen*$\rangle$ `--blmax` $\langle$*maxBranchLen*$\rangle$ `--lh-epsilon` $\langle \epsilon_{\mathrm{LnL}} \rangle$ `--param-eps` $\langle \epsilon_{\mathrm{model}} \rangle$ `--brlen-smoothings` $\langle$*num_iters*$\rangle$ `--spr-lheps` $\langle \epsilon_{\mathrm{brlen}} \rangle$ `--bfgs-factor` $\langle$*bfgs_factor*$\rangle$ `--seed` $\langle$seed$\rangle$ `--prefix` $\langle$outdir$\rangle$ |
| Search | RAxML-NG *Study 4* | `raxml-ng --msa` $\langle$MSA$\rangle$ `--model` $\langle$model/partition$\rangle$ `--tree` $\langle$pars{1}/rand{1}$\rangle$ `--blmin` $\langle$*minBranchLen*$\rangle$ `--blmax` $\langle$*maxBranchLen*$\rangle$ `--lh-epsilon-auto` $\langle \epsilon_{\mathrm{auto}} \rangle$ `--lh-epsilon-fast` $\langle \epsilon_{\mathrm{fast}} \rangle$ `--lh-epsilon-slow` $\langle \epsilon_{\mathrm{slow}} \rangle$ `--lh-epsilon-brlen-full` $\langle \epsilon_{\mathrm{brlen\_full}} \rangle$ `--param-eps` $\langle \epsilon_{\mathrm{model}} \rangle$ `--brlen-smoothings` $\langle$*num_iters*$\rangle$ `--spr-lheps` $\langle \epsilon_{\mathrm{brlen}} \rangle$ `--bfgs-factor` $\langle$*bfgs_factor*$\rangle$ `--seed` $\langle$seed$\rangle$ `--prefix` $\langle$outdir$\rangle$ |
| Search | IQ-TREE | `iqtree -s` $\langle$MSA$\rangle$ $\langle$`-m` model / `-p` partition$\rangle$ `-ninit 1` `-blmin` $\langle$*minBranchLen*$\rangle$ `-blmax` $\langle$*maxBranchLen*$\rangle$ `-me` $\langle \epsilon_{\mathrm{model}} \rangle$ `-eps` $\langle \epsilon_{\mathrm{LnL}} \rangle$ `-seed` $\langle$seed$\rangle$ `-pre` $\langle$outdir$\rangle$ |
| Search | FastTree | `fasttree -`$\langle$model$\rangle$ `-blmin` $\langle$*minBranchLen*$\rangle$ `-lheps` $\langle \epsilon_{\mathrm{LnL}} \rangle$ `-seed` $\langle$seed$\rangle$ `<` $\langle$MSA$\rangle$ `>` $\langle$outdir$\rangle$ |
| Evaluation | RAxML-NG | `raxml-ng --eval --msa` $\langle$MSA$\rangle$ `--model` $\langle$model/partition$\rangle$ `--tree` $\langle$tree$\rangle$ `--blmin` $\langle$*minBranchLen*$\rangle$ `--blmax` $\langle$*maxBranchLen*$\rangle$ `--lh-epsilon` $\langle \epsilon_{\mathrm{LnL}} \rangle$ `--param-eps` $\langle \epsilon_{\mathrm{model}} \rangle$ `--brlen-smoothings` $\langle$*num_iters*$\rangle$ `--bfgs-factor` $\langle$*bfgs_factor*$\rangle$ `--seed` 0 `--prefix` $\langle$outdir$\rangle$ |
| Evaluation | IQ-TREE | `iqtree -s` $\langle$MSA$\rangle$ $\langle$`-m` model / `-p` partition$\rangle$ `-te` $\langle$tree$\rangle$ `-blmin` $\langle$*minBranchLen*$\rangle$ `-blmax` $\langle$*maxBranchLen*$\rangle$ `-me` $\langle \epsilon_{\mathrm{model}} \rangle$ `-eps` $\langle \epsilon_{\mathrm{LnL}} \rangle$ `-seed 0 -pre` $\langle$outdir$\rangle$ |
| Significance Tests | IQ-TREE | `iqtree -s` $\langle$MSA$\rangle$ $\langle$`-m` model / `-p` partition$\rangle$ -z $\langle$all_trees$\rangle$ `-te` $\langle$best_tree$\rangle$ `-n 0 -zb 10000 -zw -au -pre` $\langle$outdir$\rangle$ |
| Search | RAxML-NG *Study 3* | `raxml-ng --search --msa` $\langle$MSA$\rangle$ `--model` $\langle$model/partition$\rangle$ `--lh-epsilon` 10 `--spr-lheps` 1000 `--seed` 0 `--prefix` $\langle$outdir$\rangle$ |
| Bootstrapping | RAxML-NG | `raxml-ng --bootstrap --msa` $\langle$MSA$\rangle$ `--model` $\langle$model/partition$\rangle$ `--lh-epsilon` $\langle \epsilon_{\mathrm{LnL}} \rangle$ `--spr-lheps` $\langle \epsilon_{\mathrm{brlen}} \rangle$ `--seed` $\langle$seed$\rangle$ `--prefix` $\langle$outdir$\rangle$ |
| Boostrap Support | RAxML-NG | `raxml-ng --support --tree` $\langle$tree$\rangle$ `--bs-trees` $\langle$boostrap replicates$\rangle$ `--prefix` $\langle$outdir$\rangle$ |
| Bootstrap Consensus | RAxML-NG | `raxml-ng --consense MRE --tree` $\langle$bootstrap replicates$\rangle$ `--prefix` $\langle$outdir$\rangle$ |

Table S2: The command lines we use to conduct tree inferences, re-evaluations, significance tests, and bootstrapping analyses with RAxML-NG, IQ-TREE, and FastTree.

IQ-TREE implements the following significance tests: the Kishino-Hasegawa test [8] and the Shimodaira-Hasegawa test [17], both in their weighted and unweighted variants, the Approximately Unbiased test [18], as well as the Expected Likelihood Weight test [24]. We use the default IQ-TREE settings for the number of resampling of estimated log-likelihoods (RELL) replicates (10 000) and the significance level ($\alpha = 0.05$). Since the significance tests can be biased by the number of trees in the candidate set [24], we remove topologically identical trees before applying the tests.

##### 1.3 Pipeline Setup

We implement the analysis pipeline as described in the main paper, using the Snakemake workflow management system [9] and Python 3. The pipeline code is available at `https://github.com/tschuelia/ml-numerical-analysis`. We execute this analysis pipeline on two institutional clusters (Cascade and Haswell) at the Heidelberg Institute for Theoretical Studies (HITS) and three stand-alone servers of our research group:

- Cascade: 150 compute nodes with Intel Cascade Lake CPUs (Intel Xeon Gold 6230). Each node has 20 cores with 2.1 GHz and 96 GB RAM.
- Haswell: 224 compute nodes with Intel Haswell CPUs (E5-2630v3). Each node has 16 cores with 2.4 GHz and 64 GB RAM.
- Group Server 1: AMD EPYC 7413, 24 cores with 2.65 GHz and 512 GB RAM.
- Group Server 2: AMD EPYC 7452, 64 cores with 2.35 GHz and 1024 GB RAM.
- Group Server 3: Xeon Platinum 8260, 48 cores with 2.4 GHz and 754 GB RAM.

For Studies 1, 2, and 4, we use a single core and two threads for all MSAs. Since the bootstrapping procedure requires an extensive amount of runtime, we use the RAxML-NG multiprocessing option and set the number of nodes and threads according to Table S3.

#### 2 Datasets

As mentioned in the main paper, in *Study 1* we first analyze 22 empirical unpartitioned DNA MSAs (*Data collection 1*). To verify the results, we analyze additional MSAs, including amino-acid (AA) and partitioned MSAs ( *Study 2, Data collection 2*). Table S4 provides an overview of the MSAs we use. A [1] indicates that we used this MSA for *Study 1*, a [2] indicates that we used this MSA for Studies 2–4. Because of the dataset size of the MSA marked by a [2∗], we do not analyze all threshold settings during Studies 2 and 4, but only

| MSA name | nodes/threads |
|---|---|
| D4, D10, D12, D15, D25, D27, D80, D125, D354 | 1/2 |
| D37 | 4/20 |
| D46 | 2/20 |
| D58 | 30/20 |
| D70 | 6/16 |
| D101 | 1/20 |
| D103 | 6/16 |
| D140 | 8/16 |
| D775 | 14/20 |
| D815 | 10/20 |
| D1288 | 4/20 |
| D1604 | 4/20 |

Table S3: Multiprocessing scheme for the RAxML-NG bootstrapping procedure.

the suggested new settings to verify our results on a large MSA. For the unpartitioned DNA MSAs, we use the general time reversible (GTR) model [25] of nucleotide substitution with four discrete $\Gamma$ rate categories to model among site rate heterogeneity. Respectively, we use the LG substitution model [11] for AA MSAs. For partitioned MSAs, we use the partition files as provided in the respective publication. All MSAs and partition files are available for download at `https://cme.h-its.org/exelixis/material/freeLunch_data.tar.gz`.

| MSA name | # Taxa / # Sites | Data type / # Partitions | Notes and Reference |
|---|---|---|---|
| D150[1] | 150 / 1269 | DNA / 1 | rRNA gene data of microsporidia [23]. |
| D218[1] | 218 / 2294 | DNA / 1 | Prokaryotic sequences from the small ribosomal unit [6]. |
| D500[1] | 500 / 1398 | DNA / 1 | rbcL gene data (often referred to as zilla dataset) [2]. |
| D714[1] | 714 / 1241 | DNA / 1 | MSA used for benchmarking ML inference tools [22]. |
| D1481[1] | 1481 / 1241 | DNA / 1 | MSA used for benchmarking ML inference tools [22]. |
| D1512[1] | 1512 / 1577 | DNA / 1 | MSA used for benchmarking ML inference tools [22]. |
| D1718[1] | 1718 / 1371 | DNA / 1 | rbcL gene data [21]. |
| D1908[1] | 1908 / 1424 | DNA / 1 | Fungal sequence data [22]. |
| D2000[1] | 2000 / 1251 | DNA / 1 | MSA used for benchmarking ML inference tools [22]. |
| D2308[1] | 2308 / 1224 | DNA / 1 | Mammalian sequence data [22]. |
| D2445[1] | 2445 / 1371 | DNA / 1 | rbcL gene data [21]. |

| | | | |
|---|---|---|---|
| D2554[1] | 2554 / 1232 | DNA / 1 | rbcL gene data [22]. |
| D3782[1] | 3782 / 1371 | DNA / 1 | rbcL gene data [21]. |
| D4869[1] | 4869 / 28 361 | DNA / 1 | SARS-CoV2 data (snapshot 05/05/2020 obtained from gisaid.org). |
| D27[1,2] | 27 / 1940 | DNA / 1 | Data of the 28s rRNA gene of a broad taxonomic diversity [7]. |
| D37[1,2] | 37 / 1 338 678 | DNA / 1 | Data of 447 nuclear genes of Eutheria [19]. |
| D46[1,2] | 46 / 239 763 | DNA / 1 | Data of 310 nuclear genes of seed plants [29]. |
| D101[1,2] | 101 / 1858 | DNA / 1 | rRNA gene data of microsporidia [23]. |
| D125[1,2] | 125 / 29 149 | DNA / 1 | Mammalian DNA sequences [22]. |
| D354[1,2] | 354 / 460 | DNA / 1 | Internal transcribed spacer (ITS) region of nuclear ribosomal DNA of Acer [5]. |
| D1288[1,2] | 1288 / 1200 | DNA / 1 | Mammalian sequence data [22]. |
| D1604[1,2] | 1604 / 1276 | DNA / 1 | MSA used for benchmarking ML inference tools [22]. |
| D4[2] | 4 / 325 | DNA / 1 | Primates DNA sequences from the release 96 of the Ensembl Compara database [30]. |
| D10[2] | 10 / 522 | DNA / 1 | Primates DNA sequences from the release 96 of the Ensembl Compara database [30]. |
| D12[2] | 12 / 199 | DNA / 1 | Primates DNA sequences from the release 96 of the Ensembl Compara database [30]. |
| D15[2] | 15 / 936 | DNA / 1 | Primates DNA sequences from the release 96 of the Ensembl Compara database [30]. |
| D25[2] | 25 / 1051 | DNA / 1 | Primates DNA sequences from the release 96 of the Ensembl Compara database [30]. |
| D80[2] | 80 / 232 | DNA / 1 | Primates DNA sequences from the release 96 of the Ensembl Compara database [30]. |
| D815[2] | 815 / 20 364 | DNA / 29 | Mitochondrial and nuclear sequence data of bats [16]. |
| D140[2] | 140 / 1104 | AA / 1 | Papillomavirus data [15]. |
| D775[2] | 775 / 4519 | AA / 1 | MSA used for benchmarking ML inference tools [22]. |

| | | | |
|---|---|---|---|
| D70[2] | 70 / 59 725 | AA / 210 | Transcriptomes of choanoflagellates, glass sponges, two demosponges, and a deep-sea cnidarian [26]. |
| D103[2] | 103 / 145 359 | AA / 11 | Expressed nuclear genes of streptophyte and plants [27]. |
| D58[2]* | 58 / 1 806 035 | AA / 1 | 4862 genes of jawed vertebrates [3]. |

Table S4: Overview of the MSAs we use for our analyses. All MSAs are empirical MSAs.

### 3 Numerical Thresholds

Depending on the tree inference tool, we examine different numerical thresholds. In Table S5 we provide the settings we explore for each threshold and to what ML inference tool they are applicable, alongside with the respective default setting.

| Threshold | Tested Settings | Inference Tools (resp. default setting) |
|---|---|---|
| *minBranchLen* | $\{10^{-10}, 10^{-9}, ..., 10^{-2}\}^*$ | RAxML-NG ($10^{-6}$)<br>IQ-TREE ($10^{-6}$)<br>FastTree ($5^{-9}$) |
| *maxBranchLen* | $\{10, 10^2\}$ | RAxML-NG ($10^2$)<br>IQ-TREE (10) |
| $\epsilon_{\text{LnL}}$ | $\{10^{-3}, 10^{-2}, ..., 10^3\}$ | RAxML-NG ($10^{-1}$)<br>IQ-TREE ($10^{-3}$)<br>FastTree ($10^{-1}$) |
| $\epsilon_{\text{model}}$ | $\{10^{-3}, 10^{-2}, 10^{-1}\}$ | RAxML-NG ($10^{-3}$)<br>IQ-TREE ($10^{-2}$) |
| $\epsilon_{\text{brlen}}$ | $\{10^{-3}, 10^{-2}, ..., 10^3\}$ | RAxML-NG ($10^{-1}$) |
| *num_iters* | $\{16, 32, 64\}$ | RAxML-NG (32) |
| *bfgs_factor* | $\{10^5, 10^7, 10^9\}$ | RAxML-NG ($10^7$) |

*For FastTree we additionally ran its default setting $5^{-9}$.

Table S5: Varied numerical thresholds with analyzed settings and applicable inference tools. The values in parentheses indicate the default setting for the respective inference tool.

### 4 Results Study 1: Influence of Numerical Thresholds on LnL scores and Runtimes

In this section, we present the results of *Study 1* on *Data collection 1*. This section is separated into two subsections:
**1. Tree inference phase:** In the first subsection, we focus on the influence of varying numerical thresholds on tree inference. Since we presented the results for $\epsilon_{\text{LnL}}$ and $\epsilon_{\text{brlen}}$ in RAxML-NG and IQ-TREE in the main paper, we omit a detailed discussion of these results here. Instead, we show that the default settings for the remaining numerical thresholds, as well as the default $\epsilon_{\text{LnL}}$ setting in FastTree are appropriate.
**2. Tree Evaluation:** In the second subsection, we focus on the influence of varying numerical thresholds on tree evaluation. We show that the LnL score and runtime remain largely unaffected by the numerical threshold settings, as long as they are within a reasonable value range. Both RAxML-NG's and IQ-TREE's default settings fall within this range.

In analogy to the main paper, we compare LnL scores in percent rather than absolute log likelihood units, since we compare LnL scores within a broad absolute likelihood value range. The LnL scores for the 22 empirical MSAs of *Data collection 1* range between approximately −6400 (D354) and −12 300 000 (D4869). All presented figures summarize the results over all MSAs of *Data collection 1*.

#### 4.1 Tree Inference

In this section, we present our results for varying the numerical threshold settings for the tree inference process. We analyze the influence on the LnL scores and runtimes of *all* presented numerical thresholds.

##### 4.1.1 Likelihood epsilon $\epsilon_{\text{LnL}}$

**RAxML-NG and IQ-TREE** We discuss the results of RAxML-NG and IQ-TREE for this threshold in greater detail in the main paper, based on the results of the broader set of MSAs (*Data collection 2*). For the sake of completeness, we show the results of our analysis for this threshold in both ML tools for *Data collection 1*: Figures S1g, S1h, S2g and S2h.

**FastTree** With FastTree, we observe worse LnL scores for $\epsilon_{\text{LnL}}$ thresholds $> 1$. For most MSAs, the trees inferred under threshold settings of $10^2$ and $10^3$ are significantly worse than the best-known tree according to the statistical tests. This could be due to the numerous heuristic and numerical shortcuts in FastTree's implementation (see Section 1.1) and the lack of a tree evaluation option. The runtimes for $\epsilon_{\text{LnL}}$ settings $\leq 1$ exhibit no substantial speedup. FastTree's default $\epsilon_{\text{LnL}}$ setting $10^{-1}$ therefore represents an adequate choice.

##### 4.1.2 Likelihood epsilon $\epsilon_{\mathbf{brlen}}$

In analogy to the $\epsilon_{\text{LnL}}$ threshold, we discuss the influence of the $\epsilon_{\text{brlen}}$ threshold in detail in the main paper based on our analyses on *Data collection 2*. For the sake of completeness, we also show the results of our analysis for this threshold for the MSAs of *Data collection 1*: Figures S1i and S1j

##### 4.1.3 MinBranchLen

We observe that *minBranchLen* settings $\geq 10^{-3}$ yield worse LnL scores for all three ML inference tools. Due to the lack of an evaluation phase in FastTree, the degradation of LnL scores is an order of magnitude more pronounced than for RAxML-NG and IQ-TREE. Depending on the tool, the runtimes follow distinct trends. We observe that the default settings for all three inference tools are well-chosen.

**RAxML-NG** The LnL scores for trees inferred with RAxML-NG worsen on average by 0.16 % for *minBranchLen* settings $\geq 10^{-3}$. All *minBranchLen* settings in the range $10^{-9}$–$10^{-3}$ yield equally good LnL scores (variances $\leq$ 0.03 %). For RAxML-NG, the LnL scores for 12 out of 22 MSAs again worsen noticeably with a *minBranchLen* setting of $10^{-10}$ (Figure S1a). The LnL scores for this setting are on average 0.1 % worse compared to *minBranchLen* $= 10^{-9}$. Similar to the LnL scores, the runtimes for *minBranchLen* settings in the range $10^{-9}$–$10^{-3}$ are approximately equal, with a slight trend towards faster tree inferences under higher *minBranchLen* settings, while the runs with *minBranchLen* $10^{-10}$ and $10^{-2}$ are slower (Figure S1b). Given these observations, we conclude that a *minBranchLen* setting of $10^{-9}$–$10^{-3}$ during the tree inference is adequate. RAxML-NG's default setting $10^{-6}$ falls into this range.

**IQ-TREE** In general, *minBranchLen* settings $\geq 10^{-3}$ result in worse LnL scores (Figure S2a). The LnL scores under the highest analyzed setting *minBranchLen* $= 10^{-2}$ are on average 0.1 % worse. In analogy to RAxML-NG, for some MSAs, we obtain slightly worse LnL scores for the *minBranchLen* setting $10^{-10}$ ($\leq$ 0.11 %). We observe a high impact of the *minBranchLen* setting on the runtimes of IQ-TREE tree inferences (Figure S2b). The runtimes of IQ-TREE increase with smaller *minBranchLen* settings. For *minBranchLen* $= 10^{-10}$ tree inferences run on average twice as long as for *minBranchLen* $= 10^{-3}$. Interestingly, the runtime also increases if *minBranchLen* is set to $10^{-2}$. These tree inferences run on average $52 \pm 6$ % slower than tree inferences with *minBranchLen* $= 10^{-3}$. Taking into account these observations, the IQ-TREE default setting $10^{-6}$ for the *minBranchLen* threshold appears to represent a 'good' trade-off between runtime and LnL scores.

**FastTree** Similar to the other ML inference tools, the LnL scores for FastTree worsen with higher *minBranchLen* settings. With FastTree, the degradation is by an order of magnitude worse than for IQ-TREE and RAxML-NG. The trees for *minBranchLen* $= 10^{-2}$ are on average 1.7 % less likely than the respective best-known tree (Figure S3a). This is most likely due to the lack of an evaluation mode that improves the LnL scores under smaller *minBranchLen* settings. We observe the highest decline of LnL scores in our analysis for FastTree with *minBranchLen* set to $10^{-2}$ on the D4869 MSA. The LnL score under this *minBranchLen* setting decreases by 525 %. For *minBranchLen* settings $\leq 10^{-5}$ the LnL scores are approximately equal (variances $\ll 0.1\,\%$). The runtime fluctuates depending on the MSA and the *minBranchLen* setting. In general, there is a trend for smaller settings to induce longer runtimes (Figure S3b). Given these observations, the default *minBranchLen* setting $5^{-9}$ appears to represent a reasonable choice.

###### 4.1.4 Remaining Thresholds

For the *maxBranchLen* threshold, we observe no influence on the LnL scores of neither RAxML-NG nor IQ-TREE, but we do observe an influence on runtimes. With both, RAxML-NG and IQ-TREE, and depending on the MSA, a different *maxBranchLen* setting yields faster runtimes on average ($\leq 16\,\%$ differences; Figures S1c, S1d, S2c and S2d).

For the threshold $\epsilon_{\text{model}}$, we observe a minor variance in LnL scores for IQ-TREE and RAxML-NG ($\leq 0.1\,\%$; Figures S1e and S2e). For RAxML-NG, depending on the MSA, a different $\epsilon_{\text{model}}$ setting appears to be the fastest setting yet with no clear trend (Figure S1f)). For IQ-TREE, we observe faster tree inferences under higher $\epsilon_{\text{model}}$ thresholds (Figure S2f). We make a similar observation regarding runtimes for RAxML-NG's threshold *num_iters*. The LnL score is not affected by the *num_iters* setting (Figures S1m and S1n). With the threshold *bfgs_factor* of RAxML-NG, we notice no influence on the LnL scores, but a trend towards faster runs for higher settings (Figures S1k and S1l).

For all these thresholds, despite their minor variations in runtimes and LnL scores, we conclude that the respective default settings in both RAxML-NG and IQ-TREE are appropriate.

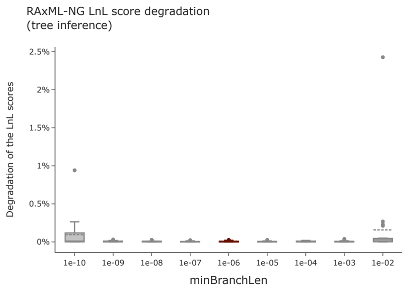


(a) Influence of the *minBranchLen* setting on the LnL scores of RAxML-NG.

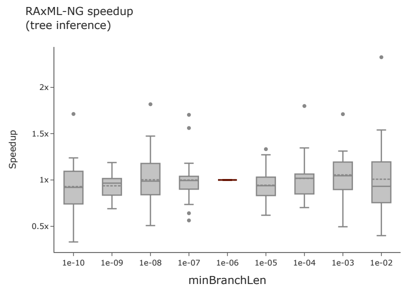


(b) Influence of the *minBranchLen* setting on the runtime of RAxML-NG.

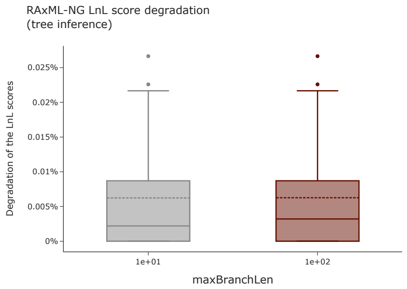


(c) Influence of the *maxBranchLen* setting on the LnL scores of RAxML-NG.

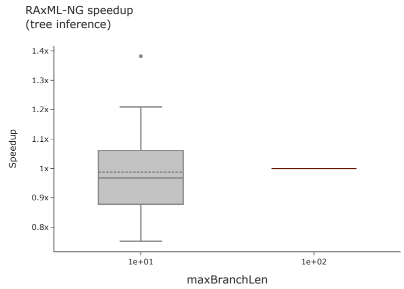


(d) Influence of the *maxBranchLen* setting on the runtime of RAxML-NG.

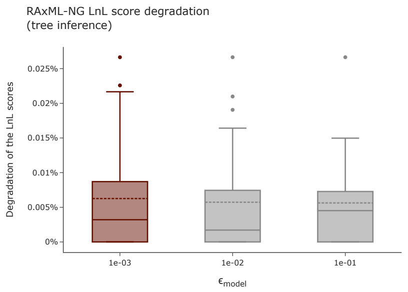


(e) Influence of the $\epsilon_{\text{model}}$ setting on the LnL scores of RAxML-NG.

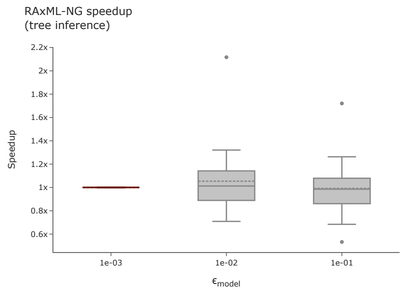


(f) Influence of the $\epsilon_{\text{model}}$ setting on the runtime of RAxML-NG.

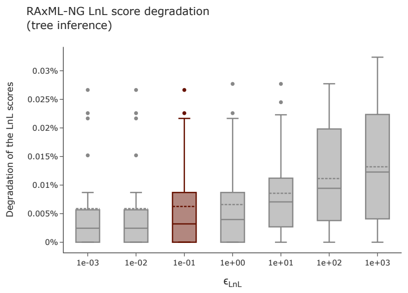


(g) Influence of the $\epsilon_{\text{LnL}}$ setting on the LnL scores of RAxML-NG.

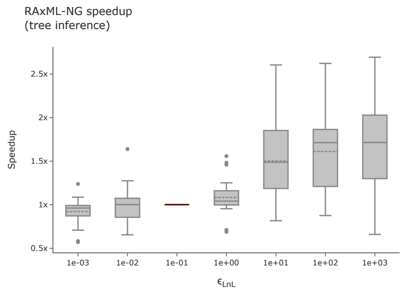


(h) Influence of the $\epsilon_{\text{LnL}}$ setting on the runtime of RAxML-NG.

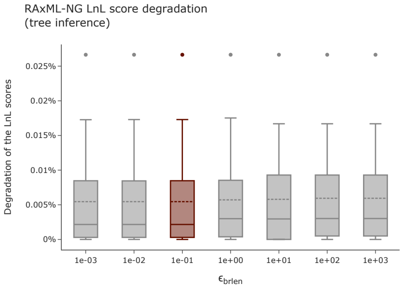


(i) Influence of the $\epsilon_{\text{brlen}}$ setting on the LnL scores of RAxML-NG.

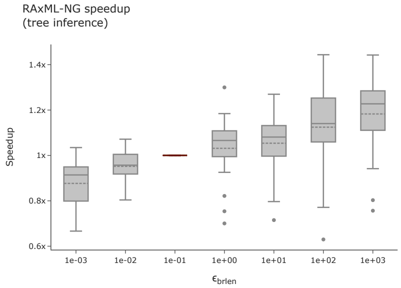


(j) Influence of the $\epsilon_{\text{brlen}}$ setting on the runtime of RAxML-NG.

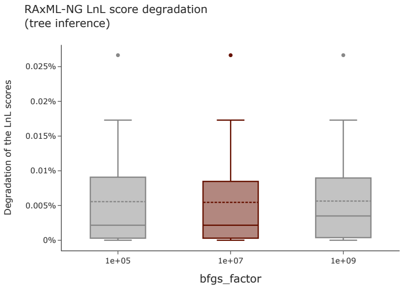


(k) Influence of the *bfgs_factor* setting on the LnL scores of RAxML-NG.

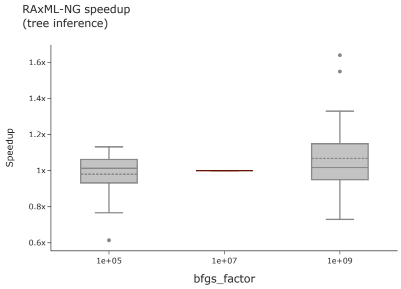


(l) Influence of the *bfgs_factor* setting on the runtime of RAxML-NG.

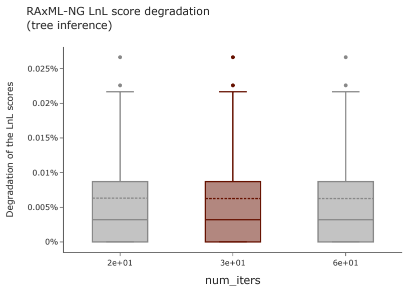


(m) Influence of the *num_iters* setting on the LnL scores of RAxML-NG.

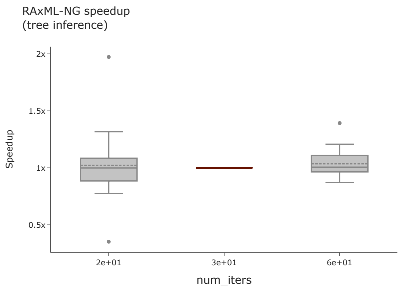


(n) Influence of the *num_iters* setting on the runtime of RAxML-NG.

Figure S1: Influence of the numerical thresholds on the LnL scores and runtimes of the RAxML-NG tree inference. For the figures in the left column, the y-axis shows the degradation in percent relative to the LnL score of the best-known tree. Higher percentages indicate worse LnL scores. For the figures in the right columns, the y-axis shows the speedup relative to the average runtime under the default setting. Each figure summarizes the data over all MSAs of *Dataset 1*. The dashed vertical line indicates the mean, and the solid vertical line the median value. The highlighted box indicates the default setting for the respective numerical threshold in RAxML-NG.

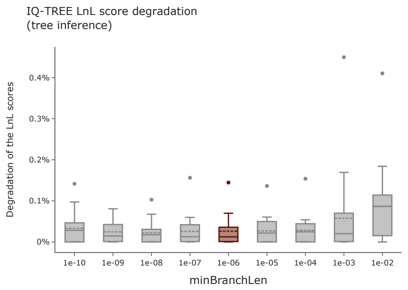


(a) Influence of the *minBranchLen* setting on the LnL scores of IQ-TREE.

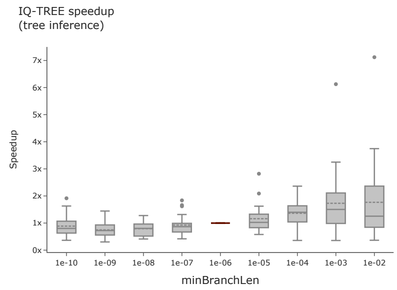


(b) Influence of the *minBranchLen* setting on the runtime of IQ-TREE.

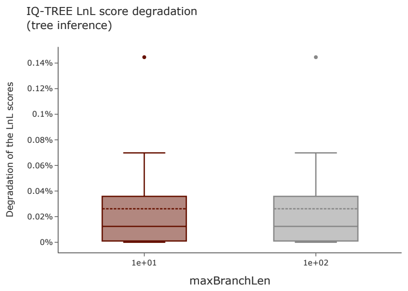


(c) Influence of the *maxBranchLen* setting on the LnL scores of IQ-TREE.

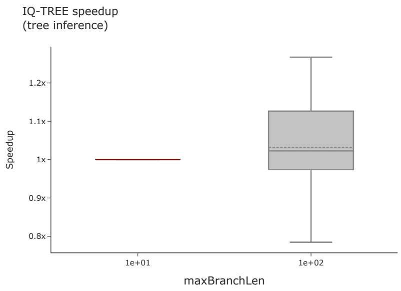


(d) Influence of the *maxBranchLen* setting on the runtime of IQ-TREE.

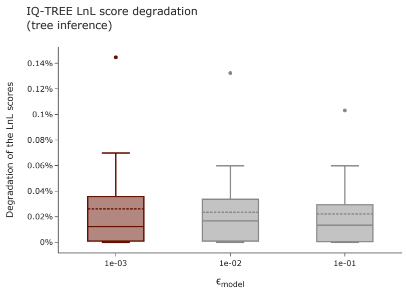


(e) Influence of the $\epsilon_{\text{model}}$ setting on the LnL scores of IQ-TREE.

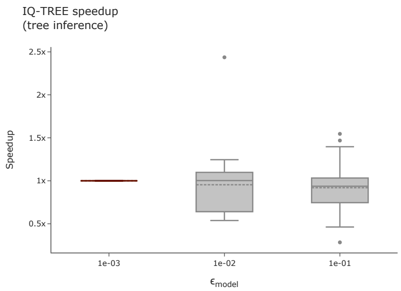


(f) Influence of the $\epsilon_{\text{model}}$ setting on the runtime of IQ-TREE.

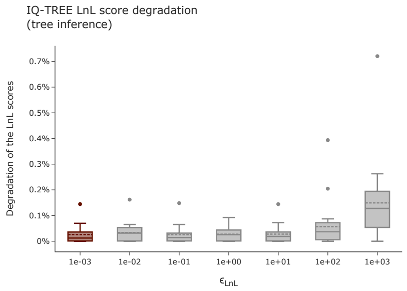


(g) Influence of the $\epsilon_{\text{LnL}}$ setting on the LnL scores of IQ-TREE.

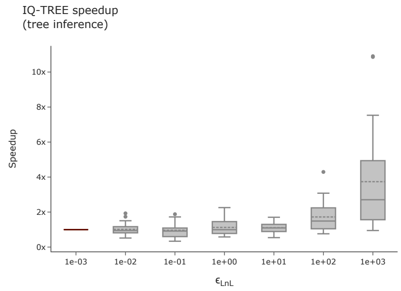


(h) Influence of the $\epsilon_{\text{LnL}}$ setting on the runtime of IQ-TREE.

Figure S2: Influence of the numerical thresholds on the LnL scores and runtimes of the IQ-TREE tree inference. For the figures in the left column, the y-axis shows the degradation in percent relative to the LnL score of the best-known tree. Higher percentages indicate worse LnL scores. For the figures in the right columns, the y-axis shows the speedup relative to the average runtime under the default setting. Each figure summarizes the data over all MSAs of *Dataset 1*. The dashed vertical line indicates the mean, and the solid vertical line the median value. The highlighted box indicates the default setting for the respective numerical threshold in IQ-TREE.

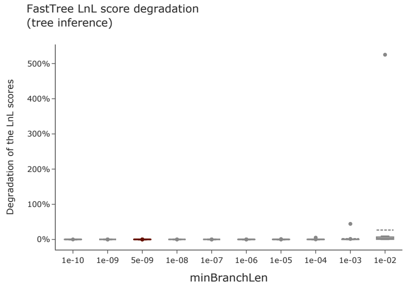


(a) Influence of the *minBranchLen* setting on the LnL scores of FastTree.

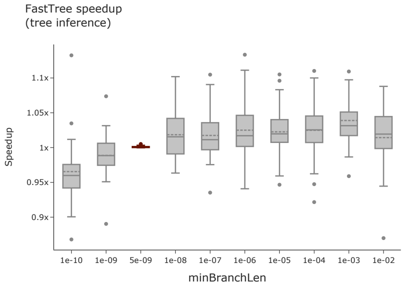


(b) Influence of the *minBranchLen* setting on the runtime of FastTree.

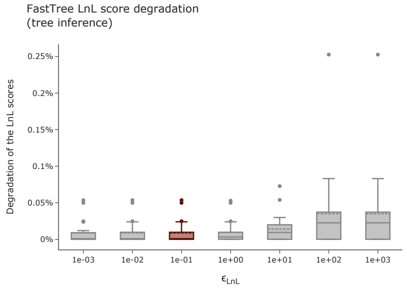


(c) Influence of the $\epsilon_{\text{LnL}}$ setting on the LnL scores of FastTree.

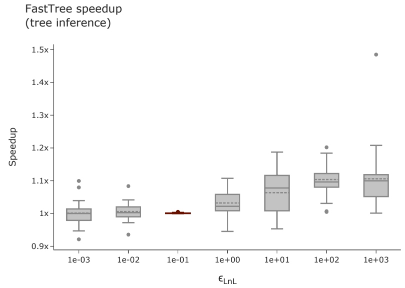


(d) Influence of the $\epsilon_{\text{LnL}}$ setting on the runtime of FastTree.

Figure S3: Influence of the numerical thresholds on the LnL scores and runtimes of the FastTree tree inference. For the figures in the left column, the y-axis shows the degradation in percent relative to the LnL score of the best-known tree. Higher percentages indicate worse LnL scores. For the figures in the right columns, the y-axis shows the speedup relative to the average runtime under the default setting. Each figure summarizes the data over all MSAs of *Dataset 1*. The dashed vertical line indicates the mean, and the solid vertical line the median value. The highlighted box indicates the default setting for the respective numerical threshold in FastTree.

#### 4.2 Tree Evaluation

In this section, we present our analysis results for varying the numerical threshold settings during tree evaluation. We analyze the influence on the LnL scores and runtimes for all considered numerical thresholds, except for the likelihood epsilon $\epsilon_{\text{brlen}}$. RAxML-NG uses this specific threshold during the SPR rounds (see Section 1.1). Since the tree topology remains unaltered during tree evaluation, this threshold is not used.

##### 4.2.1 MinBranchLen

**RAxML-NG** For RAxML-NG we observe a correlation between the LnL scores and the *minBranchLen* setting. We observe worse LnL scores with higher *minBranchLen* settings (see Figure S4a), the LnL score degradation ranges between 0.1 % and 2.0 % with a mean of 1.55 % and two outliers ((5.1 % worse for D354 and 12.5 % worse for D4869). For settings $\leq 10^{-5}$ we observe, except for D4869, equally good LnL scores. The *minBranchLen* threshold during the evaluation phase should therefore be set to $\leq 10^{-5}$. As Figure S4b shows, the runtimes for *minBranchLen* settings $\leq 10^{-5}$ are approximately identical.

**IQ-TREE** The runtimes for the IQ-TREE evaluation phase improve for *minBranchLen* settings $\geq 10^{-4}$. Averaged over all MSAs, the evaluation phase under IQ-TREE's default *minBranchLen* setting $10^{-6}$ runs $1.4 \pm 0.3$ times longer than for $10^{-2}$. For settings $\leq 10^{-4}$ we observe no clear runtime trend (Figure S5b). On 4 MSAs, we observe worse LnL scores for *minBranchLen* $\geq 10^{-3}$ ($\leq 0.3$ %; Figure S5a). We conclude that *minBranchLen* should be set to $\leq 10^{-4}$ during the IQ-TREE evaluation phase.

##### 4.2.2 Likelihood epsilon $\epsilon_{\mathbf{LnL}}$

For IQ-TREE, the likelihood threshold $\epsilon_{\mathrm{LnL}}$ does neither influence runtimes nor LnL scores (log-likelihood variances are 0.0 and runtime variances $\leq 1$ %; Figures S5g and S5h). For RAxML-NG, we observe up to 0.2 % worse LnL scores for settings $\geq 10$ and one extreme case of 1.5 % for $\epsilon_{\mathrm{LnL}} = 10^3$ (D1288; Figure S4g). Under smaller $\epsilon_{\mathrm{LnL}}$ threshold settings, we observe increased runtimes (Figure S4h). Tree re-evaluation under RAxML-NG's default $\epsilon_{\mathrm{LnL}} = 10^{-1}$ are on average 4 times slower than under $\epsilon_{\mathrm{LnL}} = 10^3$.

##### 4.2.3 Remaining Thresholds

The thresholds $\epsilon_{\mathrm{model}}$, *num_iters*, and *bfgs_factor* have no impact on the LnL score (Figures S4e, S4i and S4k). However, the runtimes for $\epsilon_{\mathrm{model}}$ increase with smaller $\epsilon_{\mathrm{model}}$ settings (on average 10.5 % increase for RAxML-NG, and 20 % for IQ-TREE; Figures S4f and S5f). The *bfgs_factor* threshold shows a similar effect: RAxML-NG runtimes increase on average 49 % with lower settings (Figure S4l). The runtimes for the *num_iters* threshold increase for more iterations (on average 3.7 %; Figure S4j). We observe no impact on, neither LnL scores, nor runtime for the *maxBranchLen* threshold in RAxML-NG (Figures S4c and S4d). For IQ-TREE we notice runtime variations $\leq 20$ %. However, depending on the MSA, a different *maxBranchLen* setting yields faster execution times (Figure S5d). The LnL score remains unaffected (Figure S5c).

Based on our findings, we suggest the following threshold settings for the evaluation phase:

| Threshold | RAxML-NG | IQ-TREE |
|---|---|---|
| *minBranchLen* | $\leq 10^{-5}$ | $\leq 10^{-4}$ |
| *maxBranchLen* | $\in \{10, 10^2\}$ | $\in \{10, 100\}$ |
| $\epsilon_{\text{LnL}}$ | $\leq 10$ | $\leq 1000$ |
| $\epsilon_{\text{model}}$ | $\leq 0.1$ | $\leq 0.1$ |
| *num_iters* | $\in \{16, 32, 64\}$ | – |
| *bfgs_factor* | $\in \{10^5, 10^7, 10^9\}$ | – |

For both tools, the respective default settings fall within our suggested value ranges. We observe that the runtime of the tree evaluation is negligible compared to the runtime of the tree inference. In our data, the average runtime of the RAxML-NG tree evaluation across all MSAs of *Data collection 1* and all numerical thresholds is $5 \pm 5\,\%$ of the runtime of the respective tree inference of a single tree. For IQ-TREE, the average runtime of the tree evaluation is $4 \pm 3\,\%$ of the runtime of the respective tree inference. Therefore, we recommend setting the numerical thresholds to their default setting during the tree evaluation.

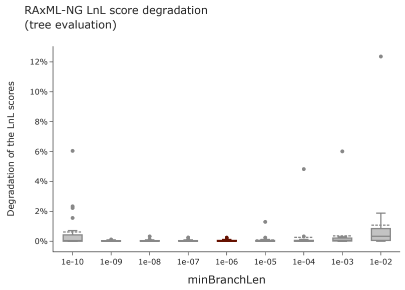


(a) Influence of the *minBranchLen* setting on the LnL scores of RAxML-NG.

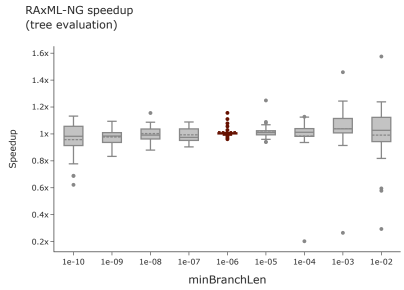


(b) Influence of the *minBranchLen* setting on the runtime of RAxML-NG.

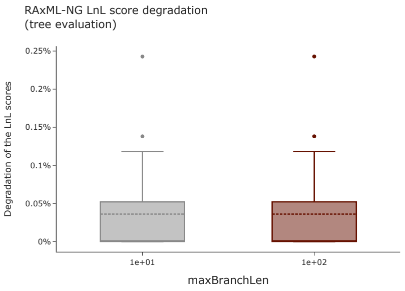


(c) Influence of the *maxBranchLen* setting on the LnL scores of RAxML-NG.

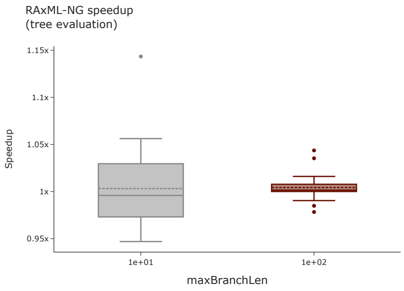


(d) Influence of the *maxBranchLen* setting on the runtime of RAxML-NG.

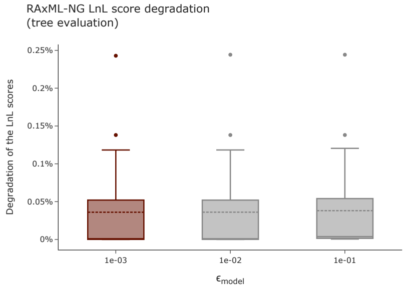


(e) Influence of the $\epsilon_{\text{model}}$ setting on the LnL scores of RAxML-NG.

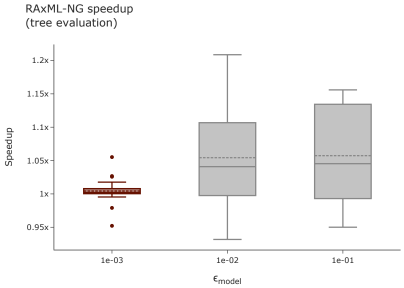


(f) Influence of the $\epsilon_{\text{model}}$ setting on the runtime of RAxML-NG.

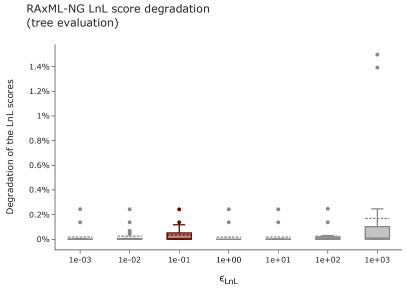


(g) Influence of the $\epsilon_{\text{LnL}}$ setting on the LnL scores of RAxML-NG.

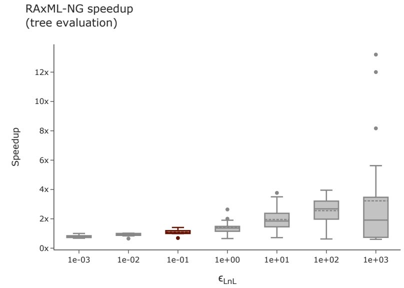


(h) Influence of the $\epsilon_{\text{LnL}}$ setting on the runtime of RAxML-NG.

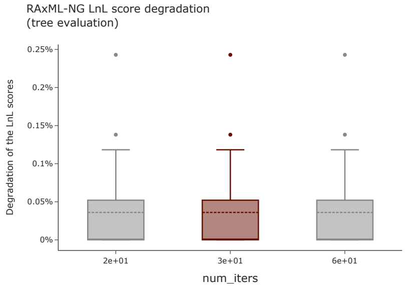


(i) Influence of the *num_iters* setting on the LnL scores of RAxML-NG.

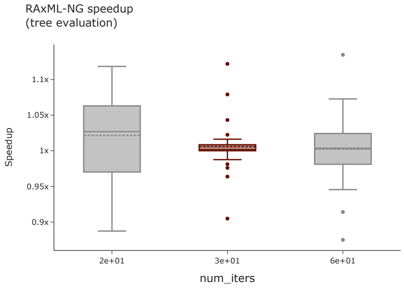


(j) Influence of the *num_iters* setting on the runtime of RAxML-NG.

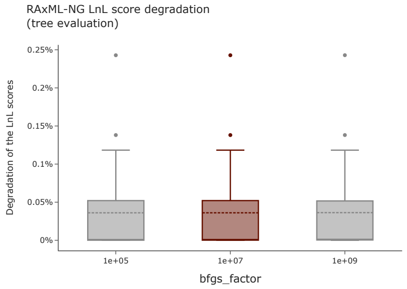


(k) Influence of the *bfgs_factor* setting on the LnL scores of RAxML-NG.

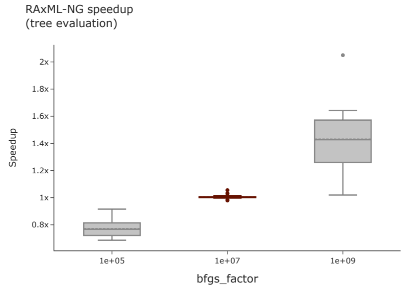


(l) Influence of the *bfgs_factor* setting on the runtime of RAxML-NG.

Figure S4: Influence of the numerical thresholds on the LnL scores and runtimes of the RAxML-NG tree evaluation. For the figures in the left column, the y-axis shows the degradation in percent relative to the LnL score of the best-known tree. Higher percentages indicate worse LnL scores. For the figures in the right columns, the y-axis shows the speedup relative to the average runtime under the default setting. Each figure summarizes the data over all MSAs of *Dataset 1*. The dashed vertical line indicates the mean, and the solid vertical line the median value. The highlighted box indicates the default setting for the respective numerical threshold in RAxML-NG.

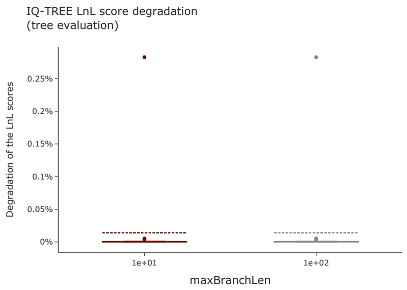


(a) Influence of the *minBranchLen* setting on the LnL scores of IQ-TREE.

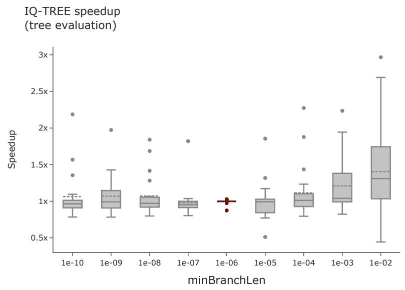


(b) Influence of the *minBranchLen* setting on the runtime of IQ-TREE.

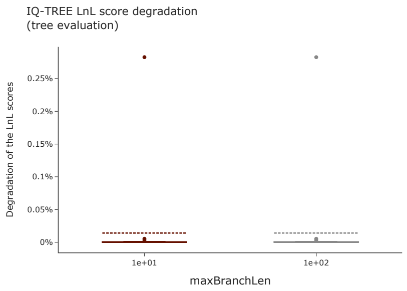


(c) Influence of the *maxBranchLen* setting on the LnL scores of IQ-TREE.

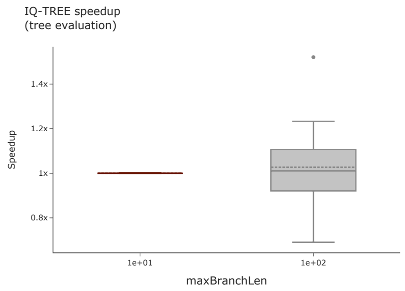


(d) Influence of the *maxBranchLen* setting on the runtime of IQ-TREE.

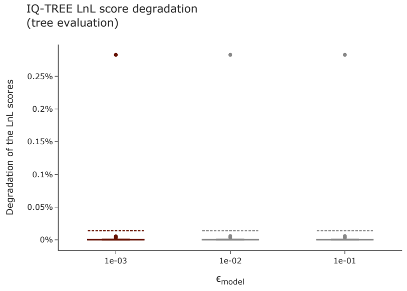


(e) Influence of the $\epsilon_{\mathrm{model}}$ setting on the LnL scores of IQ-TREE.

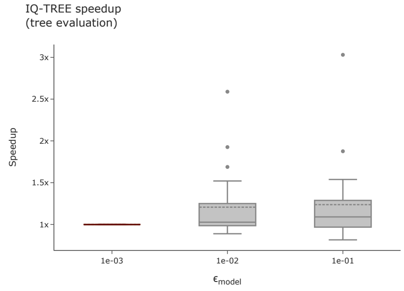


(f) Influence of the $\epsilon_{\mathrm{model}}$ setting on the runtime of IQ-TREE.

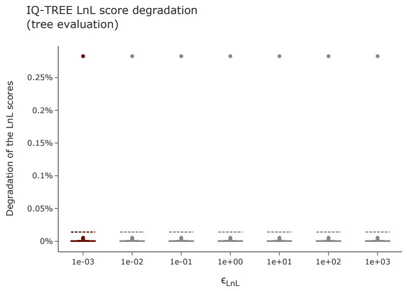


(g) Influence of the $\epsilon_{\mathrm{LnL}}$ setting on the LnL scores of IQ-TREE.

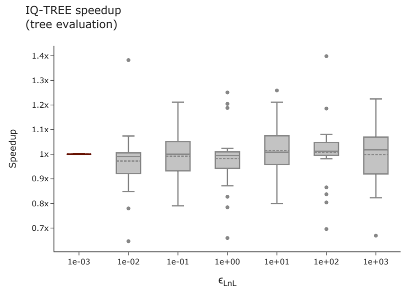


(h) Influence of the $\epsilon_{\mathrm{LnL}}$ setting on the runtime of IQ-TREE.

Figure S5: Influence of the numerical thresholds on the LnL scores and runtimes of the IQ-TREE tree evaluation. For the figures in the left column, the y-axis shows the degradation in percent relative to the LnL score of the best-known tree. Higher percentages indicate worse LnL scores. For the figures in the right columns, the y-axis shows the speedup relative to the average runtime under the default setting. Each figure summarizes the data over all MSAs of *Dataset 1*. The dashed vertical line indicates the mean, and the solid vertical line the median value. The highlighted box indicates the default setting for the respective numerical threshold in IQ-TREE.

### 5 Results Study 2: Main Paper Speedup with Outliers

In the main paper, we removed outliers for all figures depicting a speedup for Study 2. For the sake of completeness, we provide all these figures, including all outliers in Figure S6.

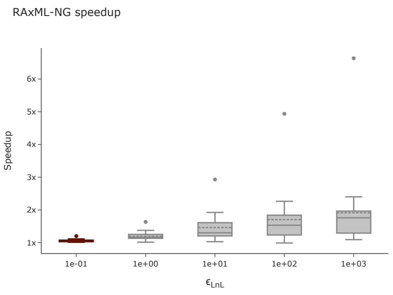


(a) Influence of the $\epsilon_{\mathrm{LnL}}$ setting on the runtime of the RAxML-NG tree inference (Corresponds to Figure 2 in the main paper).

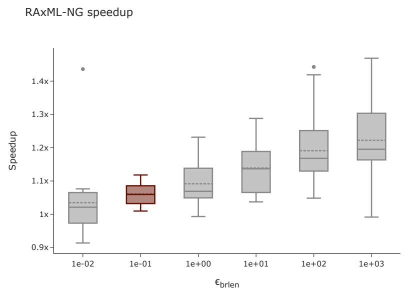


(b) Influence of the $\epsilon_{\mathrm{brlen}}$ setting on the runtime of the RAxML-NG tree inference (Corresponds to Figure 4 in the main paper).

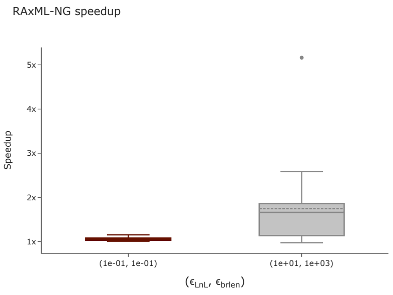


(c) Influence of simultaneously changing both likelihood epsilon settings on the runtime of the RAxML-NG tree inference (Corresponds to Figure 6 in the main paper).

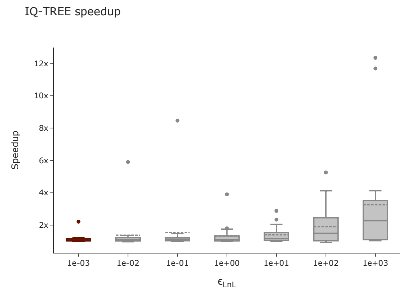


(d) Influence of the $\epsilon_{\mathrm{LnL}}$ setting on the runtime of the IQ-TREE tree inference (Corresponds to Figure 8 in the main paper).

Figure S6: These figures show the same speedup data as presented in the main paper, but include all outliers. The y-axis shows the speedup relative to the default setting of the respective ML inference tool and threshold. The dashed vertical line indicates the mean, and the solid vertical line the median value.

### 6 Results Study 4: New likelihood epsilon thresholds in RAxML-NG

As RAxML-NG uses the same threshold $\epsilon_{\text{LnL}}$ for four distinct operations during its tree inference procedure, we separate this threshold into four distinct fine-grained likelihood epsilons (see Section 1.1). The goal is to assess whether we can vary these thresholds independently and further improve upon runtime. We analyze these fine-grained threshold setting on *Data collection 2*. Our analyses show a similar behavior for all four thresholds. For the thresholds $\epsilon_{\text{auto}}$, $\epsilon_{\text{fast}}$, $\epsilon_{\text{brlen\_full}}$ we only observe a slight decrease in LnL scores under higher threshold settings (Figures S7a, S7c and S7g). With $\epsilon_{\text{slow}}$ this trend is more pronounced (Figure S7e). Our analyses show an analogous behavior for inference times. For $\epsilon_{\text{slow}}$ the runtimes decrease with higher settings (Figure S7f), for the other thresholds the runtimes are on average approximately equal under all threshold settings (Figures S7b, S7d and S7h). We conclude that a distinction does not further improve either runtime or LnL scores and is therefore unnecessary.

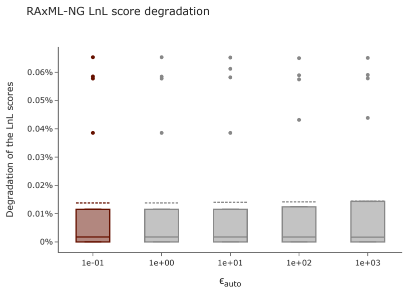


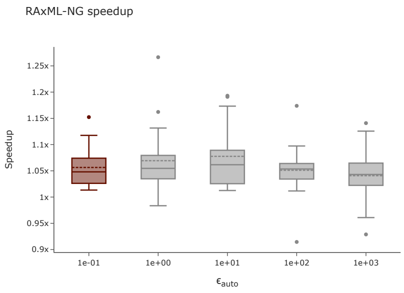


(a) Influence of the $\epsilon_{\text{auto}}$ setting on the LnL scores of RAxML-NG.

(b) Influence of the $\epsilon_{\text{auto}}$ setting on the LnL scores of RAxML-NG.

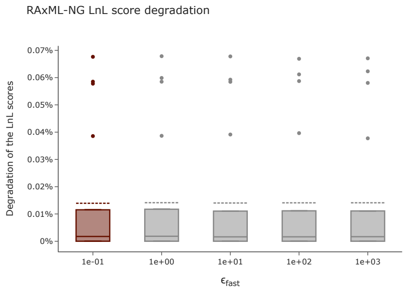


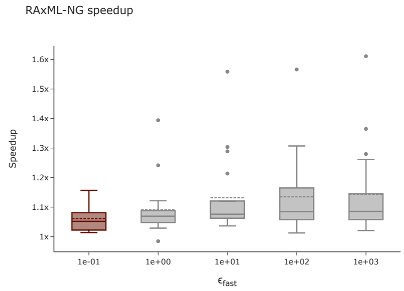


(c) Influence of the $\epsilon_{\text{fast}}$ setting on the LnL scores of RAxML-NG.

(d) Influence of the $\epsilon_{\text{fast}}$ setting on the LnL scores of RAxML-NG.

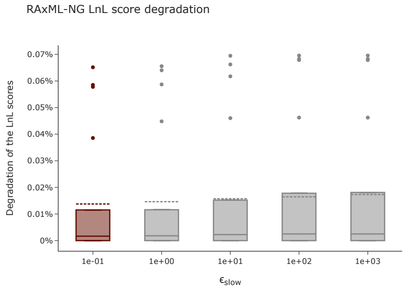


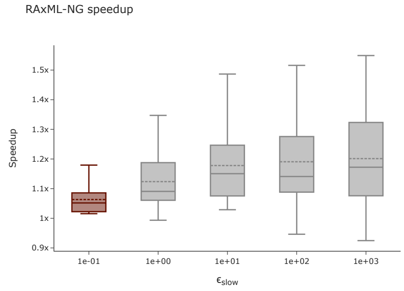


(e) Influence of the $\epsilon_{\text{slow}}$ setting on the LnL scores of RAxML-NG.

(f) Influence of the $\epsilon_{\text{slow}}$ setting on the LnL scores of RAxML-NG.

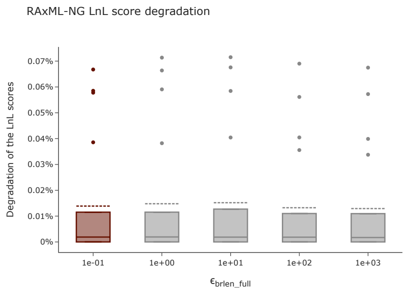


(g) Influence of the $\epsilon_{\text{brlen\_full}}$ setting on the LnL scores of RAxML-NG.

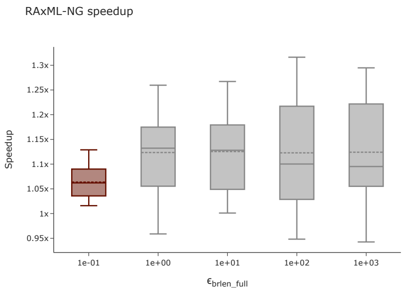


(h) Influence of the $\epsilon_{\text{brlen\_full}}$ setting on the LnL scores of RAxML-NG.

Figure S7: Influence of the separated likelihood epsilons on the LnL scores and runtimes of the RAxML-NG tree inference. For the figures in the left column, the y-axis shows the degradation in percent relative to the LnL score of the best-known tree. Higher percentages indicate worse LnL scores. For the figures in the right columns, the y-axis shows the speedup relative to the average runtime under the default setting. Each figure summarizes the data over all MSAs of *Dataset 2*. The dashed vertical line indicates the mean, and the solid vertical line the median value. The highlighted box indicates the default setting for the respective numerical threshold in RAxML-NG.

### 7 Problems with Significance Tests

As stated in the main paper, we observe problems when assessing multiple trees using statistical significance tests. Since the significance tests can be biased by the number of trees in the candidate set [24], we remove identical tree topologies from the set of inferred trees prior to applying the tests. In the following, we present an example for the rejection of trees with identical LnL scores according to the c-ELW test, as well as an example where the choice of the best tree influences the resulting significance results.

#### 7.1 c-ELW scores for identical LnL values

For trees with identical LnL values the c-ELW scores are also identical. Therefore, for trees that have a c-ELW that is close to exceeding the predefined threshold, only some trees with the exact same c-ELW score are accepted while the remaining ones are being rejected. This leads to trees being rejected despite having the same LnL score as accepted trees. Table S6 shows an example of this behavior for 6 distinct tree topologies for dataset D354. Trees 5 and 6 have identical LnL scores and identical c-ELW scores, yet only tree 5 is accepted as significant tree and tree 6 is rejected. Changing the order of trees 5 and 6 in the input file results in tree 6 being accepted, while tree 5 is rejected. Note that this issue is not due to an implementation error in IQ-TREE, but due to an unspecified behavior as per the design of the test. In addition to removing duplicate tree topologies, one would need to filter duplicate LnL values. This, however, raises the question up to how many digits LnL values are considered as being identical. This requires additional analyses, including the analysis of rounding errors and error propagation in ML inference tools that exceed the scope of our work.

| Tree ID | LnL score | c-ELW score |
|---|---|---|
| 0 | −6557.888372 | 0.254 (+) |
| 1 | −6558.638951 | 0.231 (+) |
| 2 | −6556.531087 | 0.147 (+) |
| 3 | −6556.531087 | 0.147 (+) |
| 4 | −6559.568557 | 0.142 (+) |
| **5** | **−6560.075** | **0.0393 (+)** |
| **6** | **−6560.075** | **0.0393 (-)** |

Table S6: Example of c-ELW scores for identical LnL scores. The plus sign denotes that the tree is accepted, a minus sign denotes rejection. Despite trees 5 and 6 having identical LnL scores, their tree topologies are distinct. According to the c-ELW test both trees have identical posterior weights, yet only one is accepted as plausible.

#### 7.2 Choice of the best tree

To save runtime, IQ-TREE does not re-estimate the substitution model parameters of each tree in the candidate tree set. Instead, IQ-TREE estimates these parameters using a fixed user provided tree. The choice of this tree among multiple trees with identical LnL scores influences the results of the significance tests. For example, on dataset D10 using 5 distinct tree topologies we perform the IQ-TREE significance tests 5 times, each time using a different tree as user provided tree. We observe unexpected differences between iterations for the weighted SH-Test (wSH) and the weighted KH-Test (wKH). Table S7 shows the results of this experiment. *Iter $i$* refers to the $i$-th iteration with tree $i$ used as best tree. The table shows, that during each iteration all trees have identical LnL scores, yet the significance results differ. Collecting only trees passing both tests in the plausible tree set, results in different plausible tree sets depending on the tree used as best tree.

| Test | Tree ID | Iter 0 | Iter 1 | Iter 2 | Iter 3 | Iter 4 |
|---|---|---|---|---|---|---|
| LnL | 0–4 | -124.1800658 | -124.1800886 | -124.1800431 | -124.1800886 | -124.1800431 |
| wSH | 0 | 0.0 (-) | 0.0 (-) | 1.0 (+) | 0.315 (+) | 1.0 (+) |
| | 1 | 0.0 (-) | 0.0004 (-) | 1.0 (+) | 0.739 (+) | 1.0 (+) |
| | 2 | 0.0 (-) | 0.0613 (+) | 1.0 (+) | 0.0264 (-) | 1.0 (+) |
| | 3 | 0.0 (-) | 0.86 (+) | 1.0 (+) | 0.0008 (-) | 1.0 (+) |
| | 4 | 0.0 (-) | 0.0613 (+) | 1.0 (+) | 0.0264 (-) | 1.0 (+) |
| wKH | 0 | 0.0 (-) | 0.0 (-) | 1.0 (+) | 0.0 (+) | 1.0 (+) |
| | 1 | 0.0 (-) | 0.0004 (-) | 1.0 (+) | 0.0 (+) | 1.0 (+) |
| | 2 | 0.0 (-) | 0.0 (-) | 1.0 (+) | 0.0 (-) | 1.0 (+) |
| | 3 | 0.0 (-) | 0.0 (-) | 1.0 (+) | 0.0008 (-) | 1.0 (+) |
| | 4 | 0.0 (-) | 0.0 (-) | 1.0 (+) | 0.0 (-) | 1.0 (+) |
| Plausible tree set | | {} | {} | {0, 1, 2, 3, 4} | {0, 1} | {0, 1, 2, 3, 4} |

Table S7: Influence of the choice of the best tree used to estimate the substitution model parameters on the p-values in IQ-TREE. The plus sign denotes that the tree is accepted, a minus sign denotes rejection.
